# Scale-free Vertical Tracking Microscopy: Towards Bridging Scales in Biological Oceanography

**DOI:** 10.1101/610246

**Authors:** Deepak Krishnamurthy, Hongquan Li, François Benoit du Rey, Pierre Cambournac, Adam Larson, Manu Prakash

## Abstract

Understanding key biophysical phenomena in the ocean often requires one to simultaneously focus on microscale entities, such as motile plankton and sedimenting particles, while maintaining the macroscale context of vertical transport in a highly stratified environment. This poses a conundrum: How to measure single organisms, at microscale resolution, in the lab, while allowing them to freely move hundreds of meters in the vertical direction? We present a solution in the form of a scale-free, vertical tracking microscope based on a circular “hydrodynamic-treadmill”. Our technology allows us to transcend physiological and ecological scales, tracking organisms from marine zooplankton to single-cells over vertical scales of meters while resolving microflows and behavioral processes. We demonstrate measurements of sinking particles, including marine snow as they sediment tens of meters while capturing sub-particle-scale phenomena. We also demonstrate depth-patterned virtual-reality environments for novel behavioral analyses of microscale plankton. This technique offers a new experimental paradigm in microscale ocean biophysics by combining physiological-scale imaging with free movement in an ecological-scale patterned environment.

**One sentence summary:** Scale-free vertical tracking microscopy captures, for the first time, untethered behavioral dynamics at cellular resolution for marine plankton.

---

Our oceans represent the largest habitable ecosystem on the planet. With an average depth of 4 kilometers, this unique ecosystem is highly vertically stratified with physical parameters such as light, temperature, salinity and pressure varying dramatically as a function of depth [1]. For example, only the first 200 meters of the ocean receives all the sunlight, while the deeper parts of the ocean are effectively dark. For every 10 meters in depth, the pressure increases by 1 atmosphere. Despite being only a few hundredths of the biomass of terrestrial ecosystems [2], the oceans are responsible for half of the carbon fixed on our planet [3]. Remarkably, this primary production in the ocean comes mostly from minuscule plankton [4], the majority of whom are invisible to the naked eye. Although we have known since the work of Haeckel [5] that the ocean abounds with microscopic plankton, only recently have we begun to realize their critical role in our planetary cycles [4, 6]. Despite their importance, understanding the key biophysical mechanisms at the scale of planktonic single cells and organisms, which help them navigate the ocean’s complex vertical landscape, and in turn influence planetary-scale processes, remains a major hurdle in biological oceanography.

A significant challenge in studying these biophysical processes is bridging the vast length scales (from microns to kilometers) and time scales (from millisecond to days). Conventionally, vertical fluxes in the ocean are measured using sedimentation traps and sampling at different depths [7]. Although crucial, the data is expensive to obtain and only provide a sparse window at a population level. *In situ* measurement techniques in the ocean are also advancing rapidly, including plankton imaging tools [8], and robotic tools [9], but each have certain drawbacks such as a limited tracking range and limited, macroscale resolution, respectively. In order to understand cellular physiology in this ecological context, biologists have long aspired for a technology to track single cells and organisms moving in an unbounded fluid. Here we present a framework for scale-free vertical tracking microscopy, with no bounds for motion along the vertical axis. This allows us to capture, for the first time, untethered behavioral dynamics at cellular resolution for marine plankton and other microscale objects.

## Life under gravity

Gravity and other factors such as light scattering make the ocean an extreme example of a vertically stratified ecosystem. The importance of the vertical scale in the ocean is reflected in most phytoplankton and small zooplankton having a vertically-biased motility, which is comparable to vertical flow-scales in the ocean [10, 11], while they typically passively drift in the horizontal directions. Indeed, the phenomenon of ‘Diel Vertical Migration’ (DVM), which is the largest biomass migration on our planet occurring every day, is thought to be an ecological consequence of this individual, vertically-biased motility [10, 11]. During DVM, a majority of planktonic organisms from single-celled dinoflagellates [12], and more commonly known zoo-plankton [13] rise up and sink down by several tens to hundreds of meters with a diel cycle.

Vertical movements of microscale entities relative to the ambient fluid directly affect a variety of ecological processes in the ocean. This is highlighted, in particular, by the following three classes of problems: (1) *Sedimenting particles*: Understanding the oceanic carbon sequestration process depends on the sinking rates of detritus and marine snow [14, 15], which can change due to microscopic aggregation processes and fluctuating physical parameters occurring over ecological scales. (2) *Motile plankton*: Predicting species distribution in the ocean depends on planktonic larval lifestyle of that particular species, which can be complex, involving active motility, feeding and response to cues in a vertically stratified environment [16]. (3) *Planktonic single-cells*: majority of primary production in the ocean happens in the euphotic zone and photosynthetic single-celled plankton need to regulate their depth, in many cases without the aid of motile appendages, to remain in this dynamic zone of sufficient light [17]. Therefore, in biological oceanography, the ability to measure microscale processes unravelling over ecological scales along the vertical axis, is crucial.

In the lab, concurrent observation of freely moving organisms at vastly separated scales (microns to hundreds of meters) is a fundamental experimental challenge. This is particularly true in microscopy where one has to trade-off optical resolution and field-of-view (FOV) [10]. This becomes a limiting factor in the study of objects with anisotropic motility, which rapidly move past a fixed FOV, resulting in short tracks. This issue is compounded by the fact that conventional microscopes are typically designed to image in the horizontal plane, with small chamber heights typically only a few millimeters [18]. This leads to strong interactions with the chamber walls of objects with dominant vertical movements, and a corresponding small (usable) track length. Such truncation of track lengths also leads to a statistical bias since objects with small diffusivities have a greater contribution to the tracks [10]. Tracking microscopy offers a solution to this issue where the object is kept within the optical FOV using various closed-loop tracking methods [19, 20, 21, 22, 23]. However, such approaches still limit the objects movement to the maximum size of the chamber which is ∼ 100 *mm* [22], leading to a track length that is much smaller than ecological scales (> 10 *m*). Alternative methods, such as hydrodynamic levitation using a controlled upward flow, have been used to study sedimenting marine particles [24]. However, these methods are not suitable for tracking organisms with rapidly changing behavior.

An ideal vertical tracking microscope that captures cellular details and allows unrestricted movement over ecological scales (> tens of meters) would require a meters long translational axis (Fig. 1A), and is not feasible due to cost, space and practical constraints in translating delicate optical equipment large heights. Here we present a solution to this problem, a “hydrodynamic treadmill”, that allows us to physically implement an autonomous scale-free, vertical tracking microscope. The design consists of a vertically oriented, fully-filled, circular fluidic chamber with a horizontal rotation axis (Fig. 1). A modular optical microscope is used to image the object under study with the field-of-view at either the 3 o’clock or 9 o’clock positions of the chamber, which are appropriate for compensating for a net vertical motion of the object (Fig. 1B, C). The instrument effectively provides unlimited scope for vertical motion (Fig. 1), while concurrently providing micron-and millisecond-scale spatio-temporal resolution.

**Figure 1:**
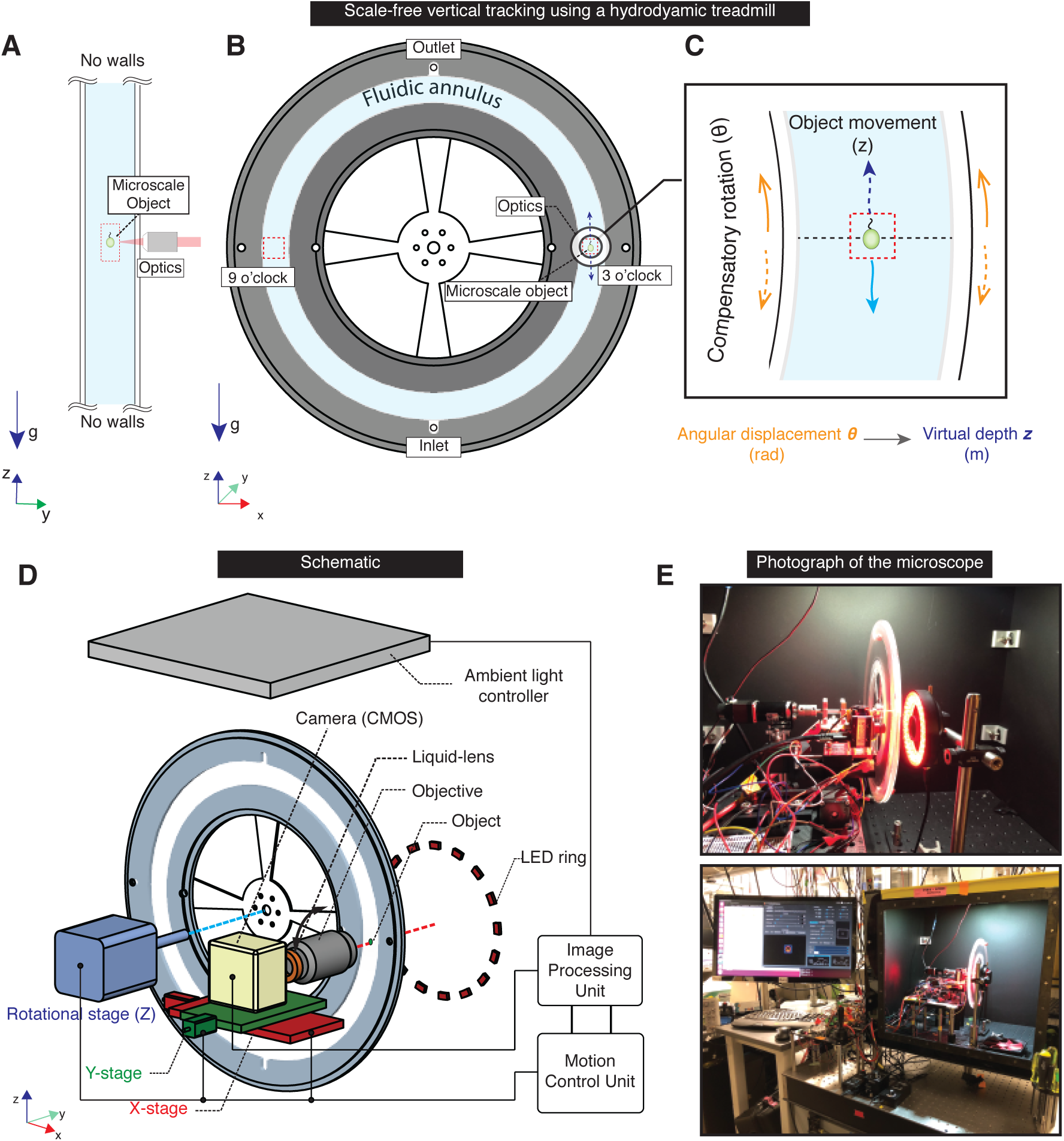
Scale-free vertical tracking microscopy using a hydrodynamic treadmill. **(A)** Schematic of an ideal, scale-free vertical tracking microscope with no boundaries in the vertical direction. **(B)** A vertically oriented circular fluidic chamber (henceforth fluidic chamber), with a contiguous annulus of fluid as a realization of such a tracking microscope. **(C)** A microscale object immersed in the ambient fluid is tracked either at the 3 o’clock or 9 o’clock positions, wherein upward or downward movements of the object relative to the fluid are compensated by stage rotation in the opposite direction using a closed-loop tracking system. Our tracking strategy, therefore, constitutes a “hydrodynamic treadmill” for microscale objects, since the vertical position of the objects is fixed in the lab reference frame. A ‘virtual-depth’ for the object can be assigned from net angular displacement and radial position. **(D)** Schematic of a practical implementation of the vertical tracking microscope showing the fluidic chamber and rotational stage for vertical (*z*) tracking, as well as translational stages coupled to the optical assembly for horizontal (*xy*) tracking. The imaging system comprises of a liquid lens assembly which is part of the focus-tracking system used to track movements along the optical axis, a CMOS sensor, and a LED dark-field illuminator. A standard desktop CPU and motion control unit (Arduino microcontroller) implement the closed-loop tracking system. **(E)** Photographs of the tracking microscope in dark-field imaging configuration with the enclosure for light and temperature control.

Previously, vertical rotating viscous flows, have been used to study the dynamics of immersed light or heavy objects [25, 26, 27], as well as for cell-culture applications in rotating-wall-vessels to simulate microgravity [28, 29]. However, such a geometry has never been used as the basis of a tracking microscope for vertically moving objects, and as such its physical feasibility, fundamental limits and design space are unknown. In order to develop this “hydrodynamic treadmill” as a scale-free vertical tracking microscope, we present a detailed theoretical analysis of the operating parameters and design space of this device. This allows us to set the operating limits and also predict the potential of our device for successfully tracking different abiotic and biotic objects.

Having characterized and calibrated the microscope, we demonstrate its capabilities by presenting several experimental vignettes drawn from the three broad classes of problems of relevance to biological oceanography, discussed above. We begin by demonstrating novel measurements of *sedimenting particles* at extreme limits of shape anisotropy, such as spheres and rods, sedimenting over macro-length and time-scales, while resolving microscale processes around the objects. Moving to particles whose physical properties are dynamically evolving due to physical and biological effects, we present Lagrangian tracks of marine detritus particles freely sedimenting over ecological length-scales of 10 meters, while concurrently observing sub-particle-scale changes.

Next we consider biotic systems with active behavior, and perform comparative, multi-scale measurements of *marine invertebrate larvae motility* and observe novel behavioral states and microscale flows, as well as their macroscale consequences. Our measurements of free-swimming plankton, clearly demonstrate a vertically biased motility that is the possible microscale underpinning of macroscale Diel Vertical Migrations. We also demonstrate environmental patterning as a function of depth, bringing the virtual reality paradigm, which is well-established in behavioral neuroscience [30] to the world of planktonic biophysics. Finally, we demonstrate tracking of *single-celled plankton* such as diatoms and dinoflagellates freely sinking and rising, capturing microscale (milliscond) behavioral fluctuations and measuring its macroscale consequences (hours).

## Scale-free vertical tracking microscopy using a “hydrodynamic treadmill”

The key components of the scale-free vertical tracking microscope are a circular fluidic chamber with a rotational axis and a horizontally laid out microscope with two-axis tracking (Fig. 1 D). This circular fluidic chamber plays a key role for any tracked object to never see boundaries in the vertical direction (*z*). The fluidic chamber consists of an annular fluidic volume with inner and outer radii *R_i_* and *R_o_*, and width *W*, so that the local cross-section is rectangular with dimensions *L* × *W*, where *L* = *R_o_* − *R_i_* (Fig. 2A). Typical values used in experiments where *R_i_* = 85 *mm*, *L* = 15, 30 *mm* and *W* = 3.2, 4, 6 *mm*. The fluidic chamber was attached to a fine rotational stage, with a horizontal rotational axis (Fig. 1D), thus allowing the chamber to be rotated with a fine angular incremental resolution (Materials and Methods). Tracking in the horizontal plane (*xy*) was achieved by translating the optical assembly which is attached to motorized linear stages (Fig. 1C, Fig. S1).

**Figure 2:**
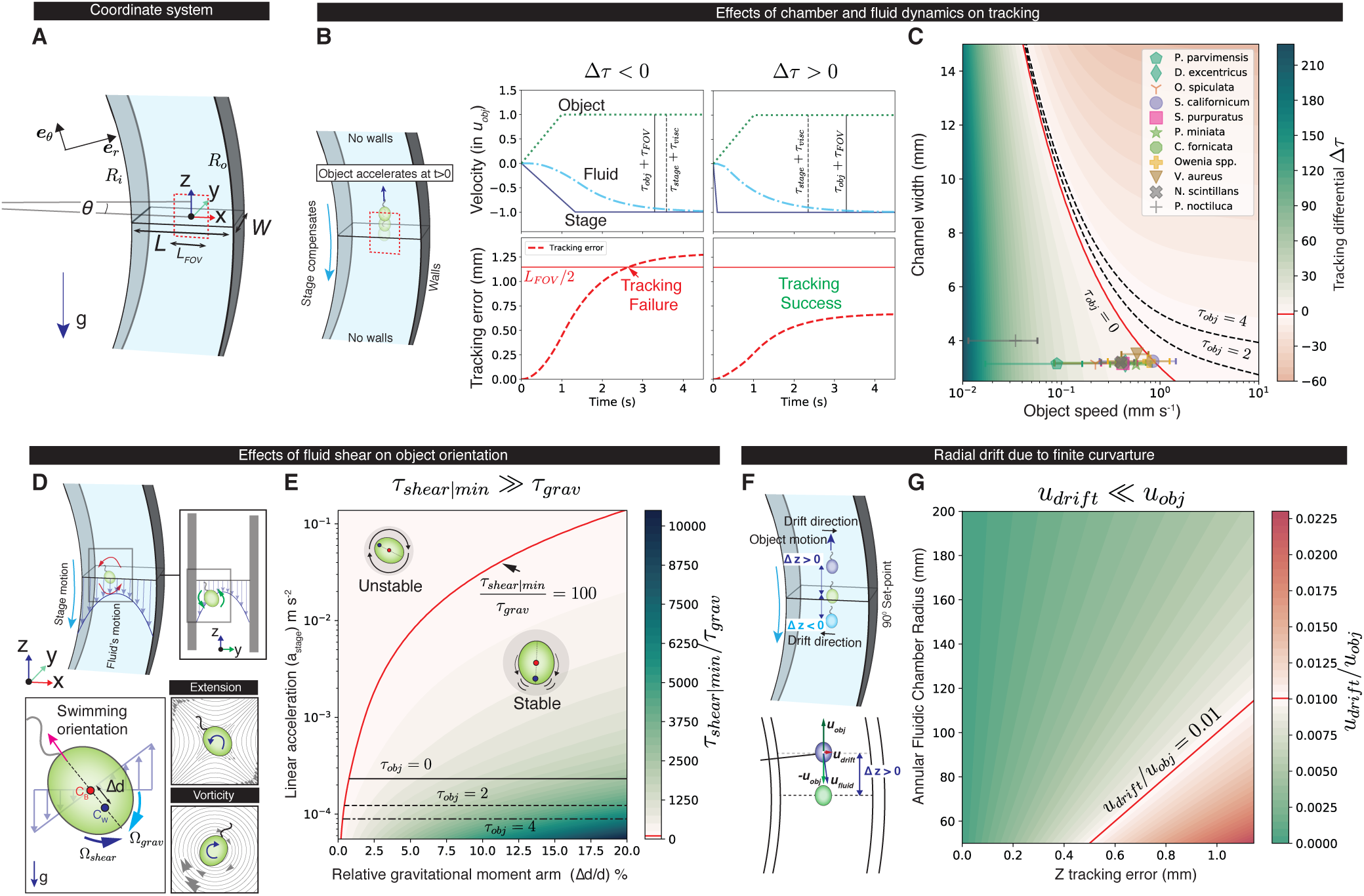
Formalism for vertical tracking using a “hydrodynamic treadmill”. **(A)** A global cylindrical coordinate system is chosen for the circular fluidic chamber, while a local cartesian coordinate system is used at the scale of the tracked object. **(B)** Left, when an object accelerates vertically to achieve a maximum velocity *u_obj_* over a behavioral time-scale *τ_obj_*, it’s motion is compensated by the stage. Velocity time traces of the object (green dashed line), stage (blue solid) and fluid (cyan dash-dotted), showing the viscous delay (*τ_visc_*) between stage and fluid movements. Tracking success can be quantified by the difference (Δ*τ*) between two sets of time-scales, one related to the object (*τ_obj_* + *τ_FOV_*) and the other related to the stage plus fluid system (*τ_stage_* + *τ_visc_*), where *τ_FOV_* = *L_FOV_ /u_obj_*, and *L_FOV_* is the optical field-of-view (FOV) size. Bottom, successful tracking is when the object’s movements can be compensated so that its distance from the center of the optical FOV (red dashed line) never exceeds half the FOV size (solid red line). **(C)** Plots of Δ*τ* with respect to the chamber width (*W*) and object speed (*u_obj_*) for *τ_obj_* = 0 (instantaneous velocity changes). The tracking limits (zero-crossing of Δ*τ*) are shown for different behavioral time-scales *τ_obj_* (solid, dashed and dot-dashed contour lines). Symbols (mean) and whiskers (standard deviation), correspond to various organisms (see Table S1) that were successfully tracked for chamber widths of 3.2, 3.5 and 4 *mm*. **(D)** Left, rotational stage acceleration leads to a non-uniform velocity profile, which at the scale of the object is locally a simple shear flow. This shear flow (with shear rate 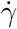), can be decomposed into an extensional and vortical component, both of which perturbs the object’s equilibrium orientation, but in opposing directions. A gravitactic effect, due to a displaced center-of-mass and center-of-buoyancy, stabilizes the orientation and aligns it with gravity over a time-scale *τ_grav_*. A balance of these two effects is quantified by the ratio of time-scales *τ_shear|min_/τ_grav_*, where *τ_shear|min_* is the inverse of the maximum possible shear rate during a chamber acceleration. **(E)** Plot of *τ_shear|min_/τ_grav_* with respect to stage acceleration and the gravitactic moment arm relative to object size, shows a broad region where the object orientation is highly stable (*τ_shear|min_/τ_grav_* ≫ 1), implying that tracking has a negligible effect on orientation. The upper bound for stage acceleration is set by imposing the condition *τ_shear|min_/τ_grav_* > 100 (solid red curve), and the lower bound is set by the condition Δ*τ* > 0 in (B) (cross-over contours shown for *τ_obj_* = 0, 2, 4). **(F)** Top, tracking a linear motion using a circular chamber implies that, for a non-zero vertical tracking error (Δ*z*), there is a radial drift induced in the object’s motion. Bottom, this drift velocity is given by *u_drift_*(*t*)*/u_obj_* = Δ*z*/*R*(*t*), where *R*(*t*) is the radial position of the object at time *t*. **(G)** Ratio of radial drift velocity to object’s speed plotted as a function of the radius of the fluidic chamber center-line ((*R_i_* + *R_o_*)/2) and the vertical tracking error, showing the cross-over (red solid line) where the ratio exceeds 1%.

We mounted a home-built light microscope focused on either the 3 o’ clock or 9 o’clock position of the circular chamber (see Fig. 1B, C; Fig. S1) such that rotational motion of the chamber resulted in tangential motion parallel to the axis of gravity, at the center of the optical field-of-view (Fig. 1C). The microscope optics consisted of a lens assembly constructed around a liquid-lens with controllable optical power (Figs. S2, S10, S11). The liquid-lens was used to rapidly modulate the focal plane of the microscope, which was used to achieve tracking along the optical axis (Supplementary Material 1.4.1, Fig. S4). This assembly was coupled to a CMOS camera, capable of full resolution color imaging at high sampling rates (238*Hz*). The modularity of the optical system allows flexibility in switching between different imaging strategies. In our experiments, we primarily used dark-field imaging using a ring LED situated on the side opposite to the imaging assembly (Fig. 1C, Fig. S1). Based on the organism being tracked, we used a suitable wavelength of dark-field illumination such that the organism’s behavior was not perturbed (Fig. S12) [31, 32].

Images captured on the camera sensor were fed to a custom image-processing pipeline implemented on a computer, and the object position was extracted rapidly (≈ 5 *ms*) using real-time machine vision algorithms (Fig. S3). The resulting tracking error, which is defined as the displacement between the object position and center of the microscope’s FOV (and focal plane), was relayed to a microcontroller, which in-turn sent motion commands to the rotational and translational stages, so as to track the object (Fig. 1D, Fig. S2). The entire experimental setup was housed within a temperature and light controlled enclosure (Fig. 1E). An isothermalization procedure was implemented such that background flows in the chamber were < 20 *µms^−1^* (Materials and Methods, Fig. S5). Ambient light intensities could be modulated as a function of ‘virtual-depth’ of the tracked object using a top-mounted white LED array so as to provide a uniform illumination across the experimental chamber (Fig. 1D,E, Fig. S1).

## A formalism for vertical tracking using a “hydrodynamic treadmill”

During tracking, small circular motions are imparted to the annulus of fluid within the circular fluidic chamber to compensate for vertical motions of the tracked object. This has two important effects: (1) Imparting rotational momentum to a contiguous annulus of fluid suffers a viscous delay leading to a transient flow profile that is not uniform (Supplementary Material 2.2, Figs. S7, S8, S9). (2) Tracking linear vertical motions using a circular motion compensation, is likely to have consequences for the measured track. The first of these is mostly a concern for biotic objects that can change their vertical velocity actively, while the second is mainly a consideration for abiotic objects with steady vertical velocities. Motivated by these unique considerations, we carried out a detailed physical analysis to identify the parameter trade-offs and design space of our vertical tracking method, in relation to the tracked object’s properties. In contrast, considerations for horizontal tracking are similar to conventional Lagrangian tracking strategies in a fluid [19, 22, 21, 20, 23], and are not discussed here. We find that a large region of parameter space exists, where microscale objects, both biotic and abiotic, can be tracked successfully, while also ensuring that the measured tracks do not differ significantly from those that would be measured in a quiescent fluid.

In general, tracked objects can undergo behavioral transitions during which their vertical velocity changes at a certain rate. We parameterized this rate using the time-scale *τ_obj_*, which is defined as *τ_obj_* = *u_obj_/a_obj_*, where *u_obj_* and *a_obj_* are the amplitudes of the velocity and acceleration changes, respectively. Compared to the response time of our vertical tracking system (*τ_tracking_*), such behavioral transitions can be slow or fast: i.e. *τ_obj_* ≫ *τ_tracking_* and *τ_obj_* ≪ *τ_tracking_*. These two limits, in general, correspond to abiotic and biotic objects, respectively. For both biotic and abiotic objects, we confirmed that for object sizes (*d* < 1 *mm*), vertical speeds (*u_obj_* < 1 *mms*^−1^) and density differentials ((*ρ_obj_* − *ρ_f_*)*/ρ_f_ ≈* 10%, where *ρ_obj_* and *ρ_f_* are the object and fluid densities) [33] relevant to marine microscale plankton and particles, the system dynamics are linear and viscous effects dominate both the object’s and fluid’s inertia (Supplementary Material 2.2). We now detail three additional physical considerations to ensure tracking success and fidelity.

### Effects of stage and fluid response times

A change in rotational speed of the circular fluidic chamber suffers a viscous time-delay (*τ_visc_*) in being transmitted to the fluid. This fluid response time scales as *τ_visc_* ∼ *W* ^2^*/ν* [34], where *W* is the chamber width (Fig. 2A, B) and *ν* is the kinematic viscosity of the fluid. Additionally, any rotational stage has a prescribed response time (*τ_stage_*) due to the specific motion-profile implemented. To quantify how these response times affect our tracking ability, we quantified the net tracking error (distance of the tracked object from the microscope’s FOV center) when an object undergoes a simulated motion, specifically, a change in vertical velocity, at a rate *τ_obj_*, by numerically, as well as analytically modelling the stage and fluid motion profiles (Fig. 2 B, center and right, Supplementary Material 2.3).

Using this model for the tracking error, we derived a tracking condition based on the constraint that the object’s tracking error should not exceed half the microscope FOV (Fig. 2B, center and right; Supplementary Material 2.3). This condition can be expressed mathematically as: *τ_stage_* + *τ_visc_* < *τ_obj_* + *τ_FOV_*, where *τ_FOV_* is the time taken for the object to swim the optical FOV size (*L_FOV_*) at its final velocity, given by *τ_FOV_* = *L_FOV_ /u_obj_*. Plotting this constraint as a tracking differential time Δ*τ* = (*τ_obj_* + *τ_FOV_*) − (*τ_stage_* + *τ_visc_*), with respect to the chamber width (*W*) and the object speed (*u_obj_*), for different behavioral transitions times (*τ_obj_*), reveals the region of parameter space where the object can be successfully tracked (Δ*τ* > 0) (Fig. 2C). We additionally show that there is a trade-off between the chamber width and object speed due to the higher viscous delays in larger chambers implying a bound on the object speeds that can be successfully tracked. Representative planktonic organisms over a wide range of sizes and swimming speeds that were tracked, lie in the successful tracking regime predicted by our analysis, hence serving as a validation (Fig. 2C, symbols). Our analysis of tracking success is conservative since we assume that objects instantaneously change their vertical velocity by their mean vertical speed, which is seldom the case.

### Effects of transient fluid motions on object orientation

Acceleration of the fluidic chamber further results in a non-uniform velocity profile, which manifests as a local simple shear flow at the scale of a microorganism (Fig. 2D, Top, Fig. S7). This shear can modify an object’s natural orientation when it exceeds gravitactic effects due to a bottom-heavy density distribution of the object [35, 36, 37] and/or shape anisotropy [38], which align the orientation of the organism with gravity (Fig. 2D left, bottom). A balance of shear and gravitactic reorientation places a limit on the maximum acceleration that can be used to track the changing vertical velocity of an object. To obtain a bound on this acceleration, we impose the condition that an object’s orientation be completely controlled by the gravitactic time scale, which for prolate spheroidal objects with a bottom heavy density distribution is given by *τ_grav_* = *µα_⊥_/*(2*g ρ_obj_* Δ*d*), where *µ* is the dynamic viscosity of the ambient fluid, *α_⊥_* is an *O*(1) viscous drag coefficient for rotation of a prolate spheroid [39], *g* is the acceleration due to gravity, and Δ*d* is the gravitactic moment arm, defined as the displacement between the center-of-buoyancy and center-of-mass of the object (Fig. 2C, bottom, left). The time scale for the shear developed is *τ_shear|min_* which is the inverse of the maximum shear rate 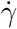*|_max_* developed during an acceleration of the fluidic chamber, and is obtained by numerically solving for the fluid flow and hence calculating the shear rate during such a motion profile (Supplementary Material 2.2). Since we consider the spatio-temporal maximum of the shear rate, which is transient, the bounds we derive, again, are conservative. The ratio of *τ_shear|min_/τ_grav_* is plotted in Fig. 2E, as a function of the maximum stage acceleration *a_stage_* and relative gravitactic moment arm Δ*d*/*d* (where the range for Δ*d*/*d* is chosen based on available measurements for microorganisms [40]). We set the condition *τ_shear|min_/τ_grav_* > 100 to obtain an upper bound for the acceleration. The lower bound is determined by the successful tracking condition, since, for very small stage accelerations, the object can easily outrun our tracking.

### Effects of finite chamber curvature

Tracking objects moving vertically using the rotational motion of the circular fluidic chamber results in a small radial drift when the vertical tracking error (Δ*z*) is finite (see Fig. 2F). This effect is most relevant for objects moving at a constant vertical velocity which is only changing slowly in time (*τ_obj_* ≫ *τ_tracking_*). We show for such an object that this drift velocity, relative to the object’s vertical velocity is given by *u_drift_/u_obj_* = Δ*z*/*R*(*t*), where *R*(*t*) is the object’s instantaneous radial position (Fig. 2F, bottom). Note that this drift is not due to a radial pressure gradient (caused by centrifugal forces) which are negligible (Supplementary Material 2.2). This ratio of velocities is plotted in Fig. 2G, as a function of the center-line radius of the annulus and vertical tracking error, and is seen to be small compared to unity for sufficiently large radius of the annulus. Thus, for most table-top setups, with *R_i_* and *R_o_* in the tens of centimeters range, this radial drift is negligible when the tracking error is within the microscope FOV which is a millimeter or smaller. For long term tracking, this error can be calculated and compensated for.

Our analysis provides a crucial design tool to optimally setup the parameters of the tracking system given some known properties of a microscale plankton or abiotic object of interest. Next, we validated our method by measuring the sedimentation speed of 250 *µm* beads of precisely calibrated density, and comparing the results to those measured using conventional Eulerian tracking in a vertical cuvette (15 *cm* tall and with same cross-section as the fluidic chamber) (Fig. S6). The two sedimentation speeds were found to have good agreement, among 6 different types of beads measured, and also found to agree with the theoretical Stokes law prediction (see Fig. S6). Having characterized and validated our tracking method, we now present several vignettes of multi-scale measurements of both abiotic and biotic systems.

## Multi-scale measurements of sedimenting particles

Sedimenting particles and their associated microscale physics are crucial to our understanding of several industrial systems and marine ecological processes [6, 14, 29, 41, 42]. In the ocean, sinking particulate organic carbon is a key driver for transport of carbon from the surface to the deep ocean, and constitutes an important component of the so called “biological pump” [14, 41]. Sinking particles are also hot-spots of biological activity in the ocean with elevated rates of photosynthesis and other biochemical processes [43, 44, 45]. Understanding tightly coupled, multi-scale processes of particle sinking, associated mass transport, and growth of biological communities are important in quantitatively understanding both microscale ecological processes on particles as well as vertical material fluxes. Motivated by this, we present four vignettes of multi-scale Lagrangian measurements of sedimenting particles in increasing order of particle complexity, wherein we concurrently measure microscale processes evolving over macroscales in length and time. We start by tracking particles whose shape is static and with different extremes of shape anisotropy (spheres and rods). We then move to particles whose shape is dynamically evolving due to flow-induced dissolution (sugar crystals). Finally, we track marine detritus particles of complex shape, whose shape, density and porosity are dynamically evolving and are controlled by both physical and biological processes.

We tracked sedimenting spheres (*d* = 500 *µm*, *ρ_obj_* = 2200 *kg m*^−3^) and observed the well-known draft-kiss-tumble interaction between pairs of spheres [46]. During this interaction the stream-wise configuration is unstable leading to the upward drafting sphere to rapidly approach the lower sphere before tumbling into a more stable cross-stream position (Fig. 3A, Left, Movie 2). Due to our tracking method, we can observe such interactions repeatedly, as seen in the time trace of sedimenting speed of one of the sedimenting spheres. Sharp peaks in the sedimentation velocity correspond to different draft-kiss-tumble events (Fig. 3A, Right).

**Figure 3:**
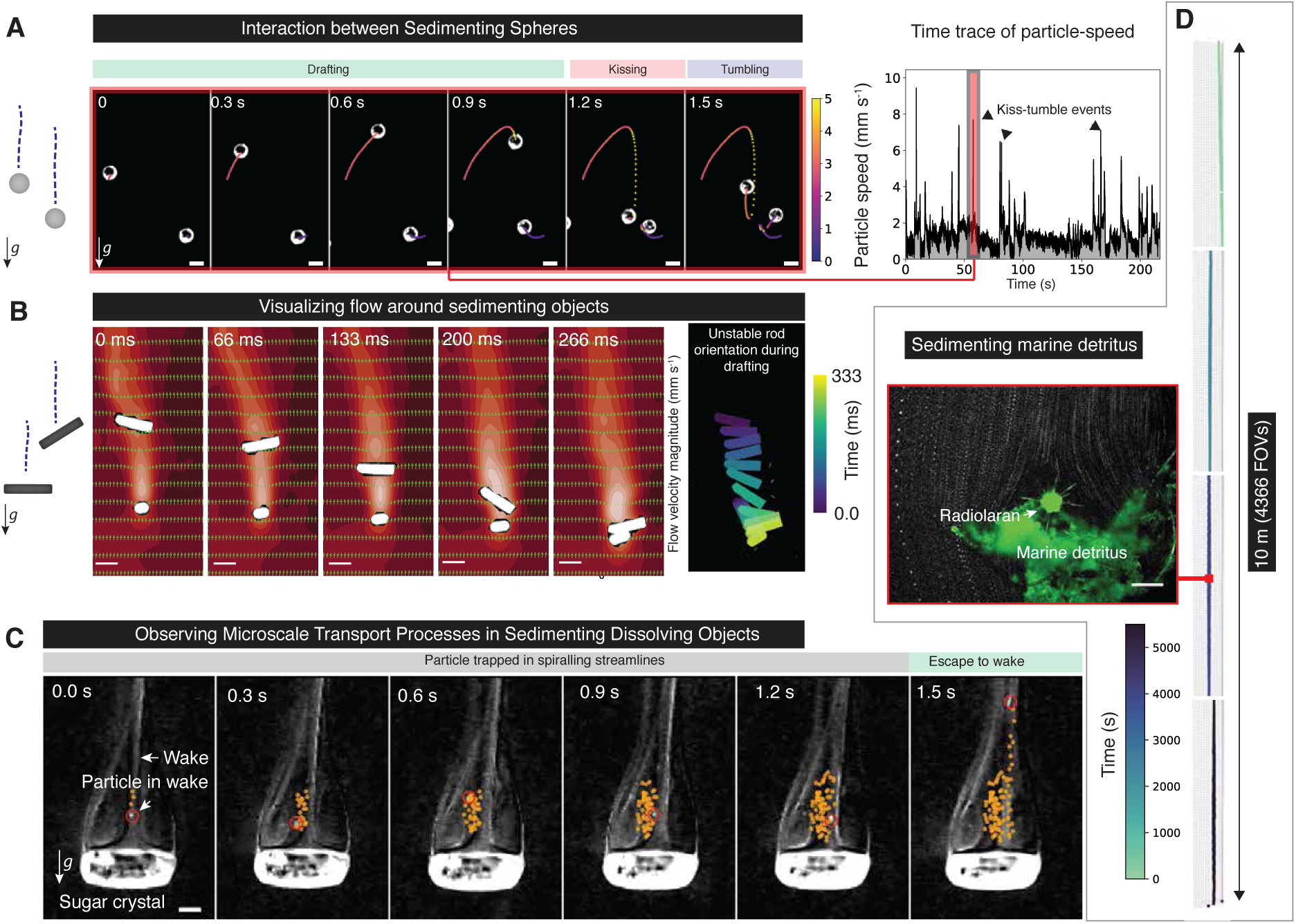
Multi-scale measurements of sedimenting particles. **(A)** Left, temporal snapshots of pair interactions between sedimenting microscale glass spheres with tracks colored by instantaneous speed. The interaction is characterized by a draft-kiss-tumble sequence (Movie 2). Scale bars, 500 *µm*. Right, long-time observations of individual spheres reveal multiple draft-kiss-tumble events which are recorded as peaks in the particle speed. **(B)** Left, temporal snapshots of flow-fields around sedimenting pairs of rods using a combination of vertical tracking, high-speed videography and Particle-Image-Velocimetry (PIV). A hydrodynamic wake behind each rod is observed. Scale bars 1 mm. Right, interaction dynamics between the rods is characterized by orientation fluctuations of the upper rod as it drafts above the lower one. **(C)** Sedimentation of a particle with dynamically changing shape due to dissolution. As an instance, we track a sugar crystal sedimenting. Temporal snapshots of particles tracked in the wake reveal a spiralling streamline region attached to the sugar crystal where particles are initially trapped before escaping to the open-streamline region at longer times. Scale bar 250 *µm*. **(D)** Tracking of sedimenting marine detritus. 3D track of the detritus sedimenting over a depth of 10 meters. Concurrent imaging of the flow around the particle reveals flow mediated material removal and rearrangements over long time-scales. The particle also contains a marine plankton (a *Radiolaran*) going along for the ride. Scale bar, 250 *µm*.

Experiments with pairs of sedimenting rods (length 2 *mm*, *d* = 500 *µm*, *ρ_obj_* = 2300 *kg m*^−3^) revealed a similar draft-kiss-tumble mechanism to that of sedimenting spheres (Fig. 3B, Left, Movie 2). Here we additionally measured the flow-field around the rods using high-speed videography and Particle-Image-Velocimetry, combined with our tracking (Fig. 3B, left). The wake behind the two rods is clearly visible and the interaction between the wakes in the streamwise configuration causes the upper rod to rapidly approach the lower rod, concurrently causing rapid orientation changes (Fig. 3B, Right).

The coupling of sedimentation and dissolution of a particle is a non-linear problem since it involves a solid interface which is dynamic, making finding analytical solutions challenging [47, 48, 49]. We present a system where this coupled process can be measured experimentally, via long-time Lagrangian tracking of sedimenting, dissolving particles. By tracking individual sedimenting sugar crystals (*d* ≈ 1 *mm*, *ρ_obj_* = 1500 *kg m*^−3^), we observed a momentum wake above the particle, consisting of a pair of recirculating regions (Fig. 3C). A thinner solute plume extends farther away from the particle (Fig. 3C, time series). As a means to measure the flow structure near the particle, we also tracked individual particles that were convected by the flow in the wake. The particles tracks revealed a spiralling streamline region adjacent to the particle, and an open-streamline region farther away (Fig. 3C, overlaid tracks, Movie 2). The presented method opens up a new avenue for studying particle dissolution in the context of sedimentation.

A related and important problem in marine ecology is observation and quantitative measurement of long distance settling of marine detritus and its associated mass transport. Using our method, we performed Lagrangian tracks of marine detritus particles collected from Monterey Bay, California (Supplementary methods). We tracked such particles as they freely sedimented over ecological length-scales of 10 *m* at a rate of *u_s_* = 2.07 ± 0.25 *mms*^−1^ (Fig. 3D). The pathlines of the flow around the detritus are shown in Fig. 3D, Bottom. Over long time scales the flow leads to particles being stripped away from the detritus, as well as rearrangement of its constituents (Movie 3). Such observations allow for the first time the direct visualization of microscale processes around a sedimenting ecosystem of high significance to global bio-geochemistry.

## Free-swimming behavior and microscale flow-fields of invertebrate larvae

Most marine invertebrate larvae have a microscale planktonic phase with complex morphology and behavior. These behaviors, which includes swimming, feeding and ultimate settlement responses are crucial in understanding the distribution of benthic ecosystems [11, 50]. Albeit its importance, it has been impossible to study this problem due to its multi-scale nature. Utilizing a diversity of species with equally diverse swimming strategies, we now demonstrate long distance tracking of free-swimming invertebrate larvae. In addition to the macroscale 3D track, we concurrently measure larval shape/posture, orientation, specific feeding and swimming behavior and hydrodynamic flow-fields. Here we present data for 8 species of marine benthic invertebrates, indigenous to the coast of Northern California, USA (Table S1). The larvae spanned 4 *Phyla* and 5 *Classes* across the animal kingdom (Fig. 4).

**Figure 4:**
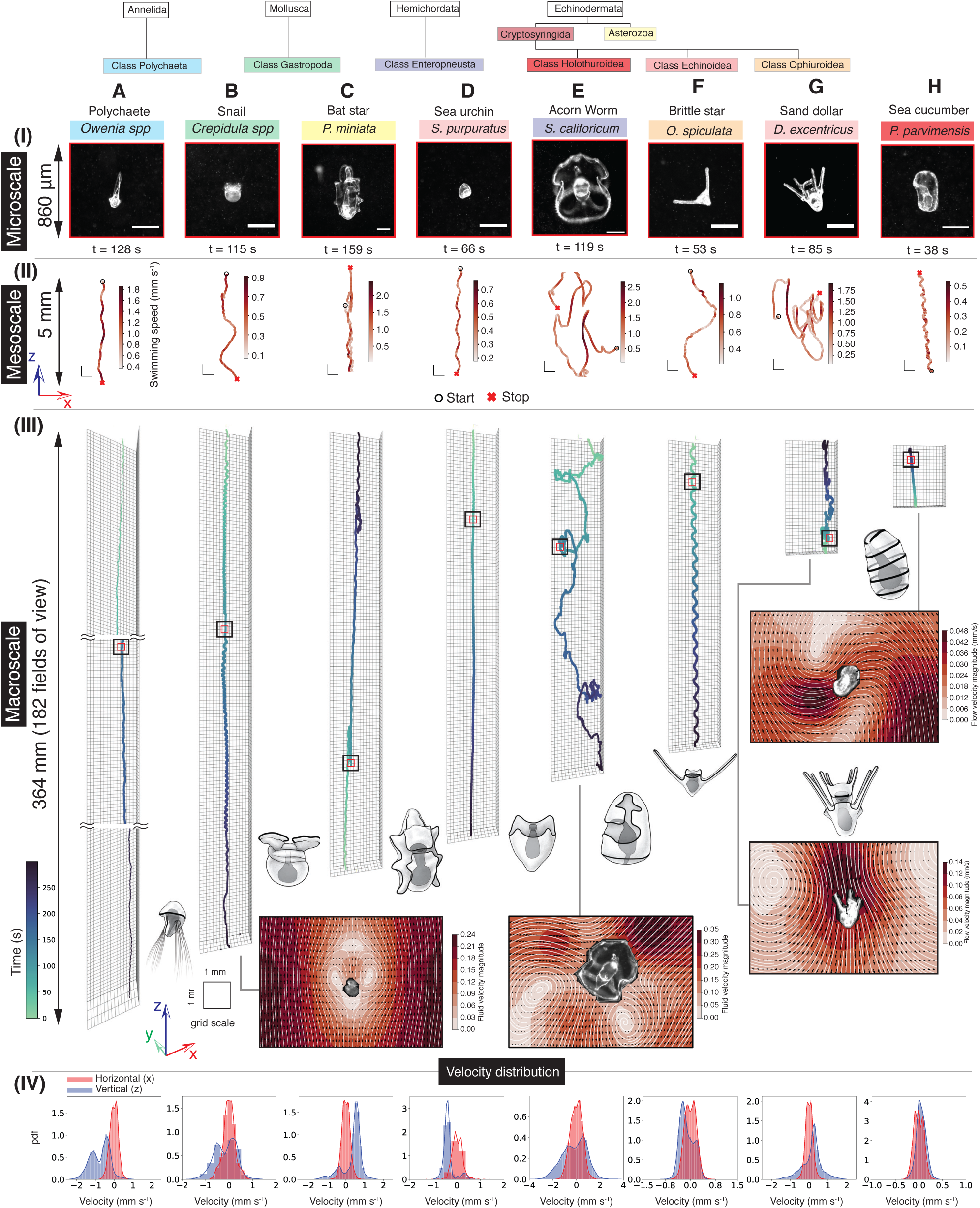
Multi-scale measurements of free-swimming behavior and flow-fields of invertebrate larvae. Columns **(A-H)**: different species of larvae. **(A)** *Owenia sp* (Poly-chaete worm), **(B)** *Crepidula sp.* (Snail), **(C)** *P. miniata* (Bat Star), **(D)** *S. purpuratus* (Purple Sea Urchin), **(E)** *S. californicum* (Acorn worm), **(F)** *O. spiculata* (Spiny Brittle Star), **(G)** *D. excentricus* (Pacific Sand Dollar) and **(H)** *P. parvimensis* (Warty Sea Cucumber). Rows **(I-IV)** Concurrent measurements of larval behavior over different length and time scales. Upper to lower rows: Increasing length and time scale. **(I)** Snapshots of freely swimming larvae from the tracks. Scale bars, 250 *µm*. **(II)** Mesoscale *xz* projection of tracks shown over a vertical extent of 5 *mm*. Scale bars, 500 *µm*. **(III)** Macroscale 3D tracks colored by elapsed track time. All track durations are truncated at 300 *s* to show the behavioral diversity of different larvae species over the same time-scale. The red (solid) and black (dashed) boxes correspond to the snapshots and mesoscale tracks shown in rows (I) and (II), respectively. The grid size is 1 *mm* × 1 *mm*. **(IV)** Distributions of vertical (*z*) and lateral (*x*) velocities for each larva, calculated over several tracks. From (III) and (IV) larvae tracked here are observed to have an anisotropic, vertically biased motility. Additionally, flow-fields around freely swimming larvae can be simultaneously measured using Particle-Image-Velocimetry. These velocity fields, in the lab frame of reference, are shown for (B), (E), (G) and (H).

We measured 3D tracks of individual larvae over time-scales of several hours and swimming over vertical extents of a few meters. Fig. 4 shows representative sub-sample of the eight larvae species over a 5 minute interval showing a diversity of swimming behaviors across scales (Movie 4). Surprisingly, all larvae exhibit vertically biased motility as seen from the anisotropic distributions of vertical and horizontal velocities (Fig. 4 (IV), Fig. S13) [4, 51]. The most striking example of this vertically biased motility was in polychaete larvae (*Owenia sp.*, Fig. 4A (III, IV)) which swam downwards a distance of 364 *mm* (182 microscope fields-of-view) over 5 minutes while having a negligible horizontal excursion (Fig. S13). At the largest scales, the vertical motility of larvae was split between active upwards or downwards swimming, where the anterior-posterior axis of the larvae was parallel or anti-parallel, respectively, to gravity; and passive/active sinking, where the larvae had an upward orientation but downward vertical velocity (Fig. 4(III), Movie 4). The multi-scale nature of the measurements revealed characteristic behaviors even at an order-of-magnitude smaller scale, as seen from the mesoscale tracks plotted over a vertical 5 *mm* extent (Fig. 4 (II)). Behavior at the mesoscale was characterized by swimming motifs including helical swimming (Fig. 4(II)A, D, F, H) free-fall (Fig. 4(II)B, C), hovering (Fig. 4(II)B, C and H) and pauses and reversals due to changes in ciliary beat (Fig. 4(II)C and G; Movie 4).

Together with tracking, we observed the microscale behavior, posture and shape of the larvae from images acquired at millisecond time-resolution (Fig. 4 (I)). By seeding the fluid with tracer particles, we measured the dynamic flow-fields around freely-swimming larvae using Particle-Image-Velocimetry (Fig. 4 I, J, K, L; Supplementary Methods). These flows, shown in the lab-reference-frame by subtracting the translational velocity of the larvae, arise from the interplay of the ciliary band-generated fluid stresses, larval shape and density (Fig. 4: Larvae sketches, Ciliary bands depicted as dark contours) [50, 51, 52, 53, 54]. In particular, we observed characteristic flows when the larvae maintained their position in the water column and created a feeding current to scan the water for food particles (Fig. 4I, L). For the larvae of *S. californicum*, we observed a characteristic dipolar velocity field due to the wide separation between the ciliary band responsible for propulsion (circular skirt at the posterior end of the larvae), and the rest of the larval body which has no propulsive contribution (Fig. 4J) [55]. In all larvae, subtracting the translational component of the velocity field revealed a strong Stokeslet contribution [22], which occurs since these larvae have an excess density to sea-water and need to create propulsive stresses just to maintain their position in the water column (Fig. 4K).

We now demonstrate that behavior of single larvae can show multi-scale dynamics. We zoom in on *P. miniata* (Bat-star) larva to further demonstrate the utility of collecting long term tracks. We measured the free-swimming behavior of these larvae (10 larvae, total track duration ≈ 1.5 hours) and found a vertically biased motility where the larvae swam upwards in a helical path (Fig. 5A) with an associated rotation of the larva body (Movie 5). We observed that the upward swimming was punctuated by a rapid behavioral transition (henceforth refered as ‘blink’) from upward to downward movement (Fig. 5 B, *z*-track). Previously, these blinks have been discussed in the context of neuronal control of ciliary band in tethered larvae [56]. Using long time scale tracks, we report the distribution and repeatable nature of this blinking behavior in free-swiming larvae. At the macroscale, blinks lead to the larvae sinking for brief periods thereby affecting their depth in the water column (Fig. 5B, z-track). Upon microscale interrogation of the blinks, we found that the blinks consisted of two behavioral micro-states: a period of passive sinking followed by a period of hovering/slower-sinking. We also found that dynamic changes in the flow-field around the larva accompanied these behavioral changes. During upward swimming we measured a streaming flow past the larvae (Fig. 5F, 1), however during the hovering/slow-sinking phase a striking array of up to four recirculating regions of flow attached to the larval body were found to develop (Fig. 5F, 2,3,4). These flows, measured for the first time in untethered organisms, are part of the larva’s feeding behavior by allowing it to stay in a relatively fixed location in the water column, using gravity as a tether, while scanning a volume of water for food particles [50, 51, 54, 57, 58, 59].

**Figure 5:**
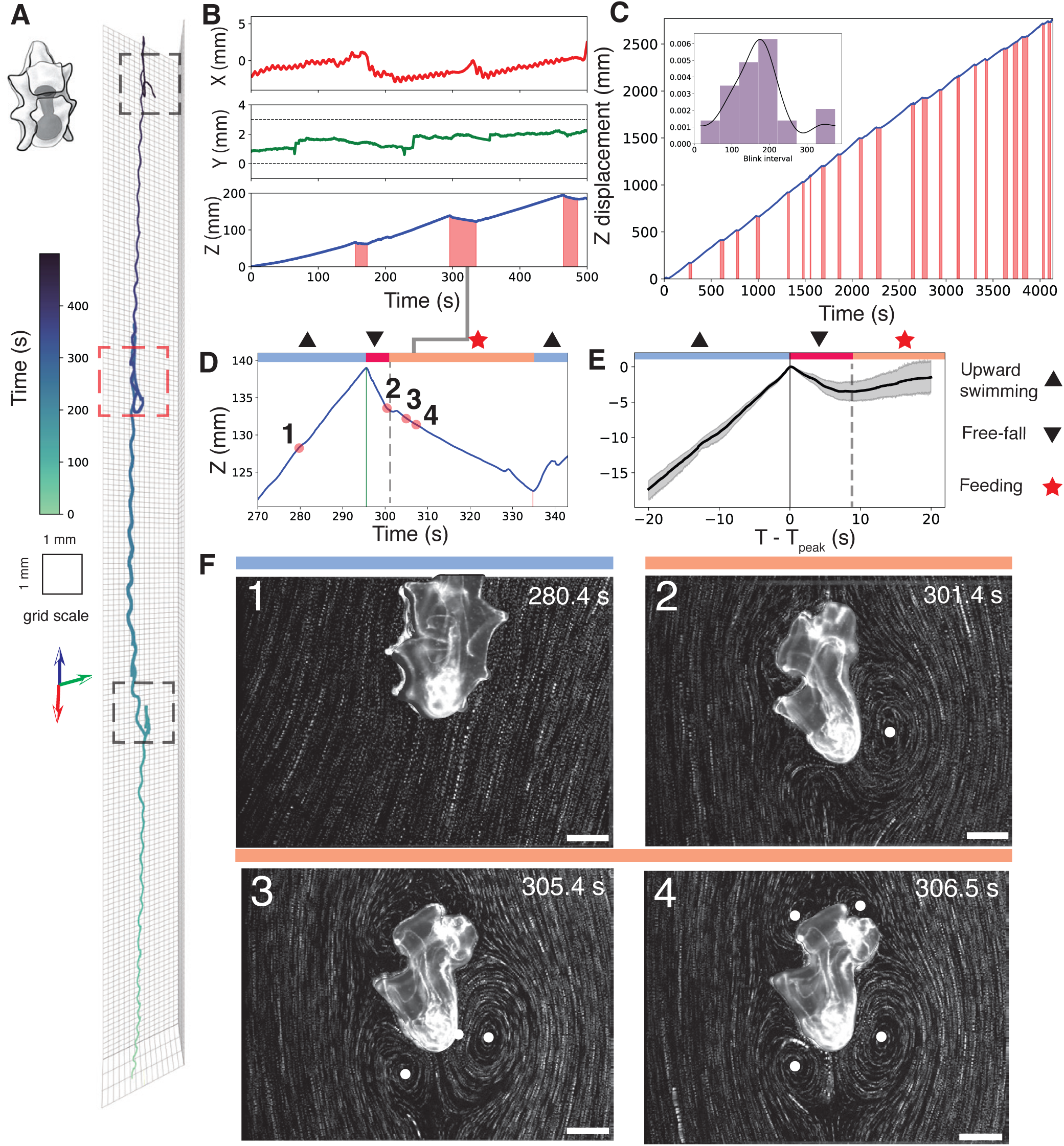
Multi-scale behavioral measurements of *P. miniata* larvae. **(A)** Three-dimensional track showing the larva swimming upwards in a columnar trajectory punctuated by three behavioral transitions (dashed boxes), henceforth “blinks” (Movie 5). **(B)** Position time-traces reveals that these blinks involve a transition from upward swimming to periods of downward motion (red bands, bottom). **(C)** Another track showing the vertical displacement of single larvae over much longer times (> 1 *hour*) with 23 behavioral transitions (red bands) while the larva swims up a distance of ≈ 2500 *mm* (≈ 1250 microscope fields-of-view). Inset, the distribution of the interval between blinks with mean and standard deviation (175 ± 78 *s*). **(D)** The high spatio-temporal resolution tracking uncovers *three* distinct behavioral sub-states during each blink consisting of a sequence of upward-swimming (upward triangle), free-fall (downward triangle), feeding (star) and upward swimming. **(E)** Behavior averaged by centering the tracks at the transition point from upward to downward displacement, over 23 blinks (mean, solid line and 95% confidence interval, gray filled region) shows the typical character of this behavioral sequence. **(F) (1-4)** Snapshots of fluid path-lines around the larvae during a blink show transitions in the flow-field as a function of the behavioral sub-states. During upward swimming the flow has the signature open-streamlines as seen in **(F)(1)**, whereas in the feeding state the larva creates up to *4* recirculating regions (centers marked by white dots) adjacent to its body (Movie 5). Feeding also involves a characteristic change in larval body shape and a widening of the oral hood (**(F) (2-4)**). Scale bars, 250 *µm*.

Our tracking method allowed us to follow individual larvae over long times (Fig. 5C, track duration ≈ 4000 *s*), which allowed us to robustly measure the statistics of these behavioral transitions, which lasted 28 ± 13 *s* and occurred with a time interval 175 ± 78 *s* (Fig. 5C, inset). Long time measurements also allowed us to extract the sterotypical behavior at the microscale during these transitions by aligning the vertical displacement tracks at the transition point from upward swimming to sinking (Fig. 5E). The behavior shows a universal profile where the sequence of micro-behavioral states of upward-swimming, sinking, feeding, followed by upward-swimming again, were conserved across a large number of blinks (Fig. 5 E, *n* = 23). Our multi-scale measurements highlight the dual function of blinks, involving macroscale (depth regulation) [60] and microscale (feeding) [50, 51, 54, 57, 58, 59] components that are intimately linked.

## Diel migration at the scale of individual plankton

An important macroscale consequence of plankton behavior is diel vertical migration where plankton rise towards the surface during night and return to deeper waters during the day. How organisms regulate this diel behavior, and hence their position in the water column as a function of different environmental cues is an area of active research [32, 60, 61]. Although the phenomena is deeply linked to circadian rhythms over a 24hr cycle, previous experimental efforts have focused on the behavior of populations [32] or have been technically limited in accessing the behavior of single organisms or individual cells over long time scales. This has led to challenges in reconciling the extent of vertical migrations observed in the field to lab measurements of motility [10].

We measured vertical migrating behavior of Polychaete larvae (*Owenia sp.*) during different times of the day while maintaining other experimental conditions constant. In particular, the larvae were imaged in red light (peak spectrum 620 nm, Fig. S12), to which they are insensitive [32] (so as to not provide any light cues) and further in a medium with no food sources, thereby representing a low-information environment. Tracks of different larvae (*n* = 6, total track duration 616 *s*) measured during day-time hours (between 1 PM and 3 PM, local time) showed a markedly downward vertical swimming velocity (*v_z_* = −0.92 ± 0.57*mms*^−1^), while those measured at night (between 8 30 PM and 9 PM, local time) swam upwards (*v_z_* = 0.42 ± 0.44*mms*^−1^, *n* = 5, total track duration 360 *s* (Fig. 6A, B). Interestingly, this swimming was active in both directions with larvae reorienting so as to point upwards or downwards to achieve swimming in that direction (Fig. 6A, inset; Movie 6). Such measurements of active upwards and downwards swimming have been reported for planktonic larvae at the scale of populations [32], but not at the scale of individual organisms. The capability to observe long distance diel migration of a freely swimming single organism, at sub-cellular resolution, opens up the capacity to unravel how internal circadian regulation [62] and external cues program this fascinating behavior.

**Figure 6:**
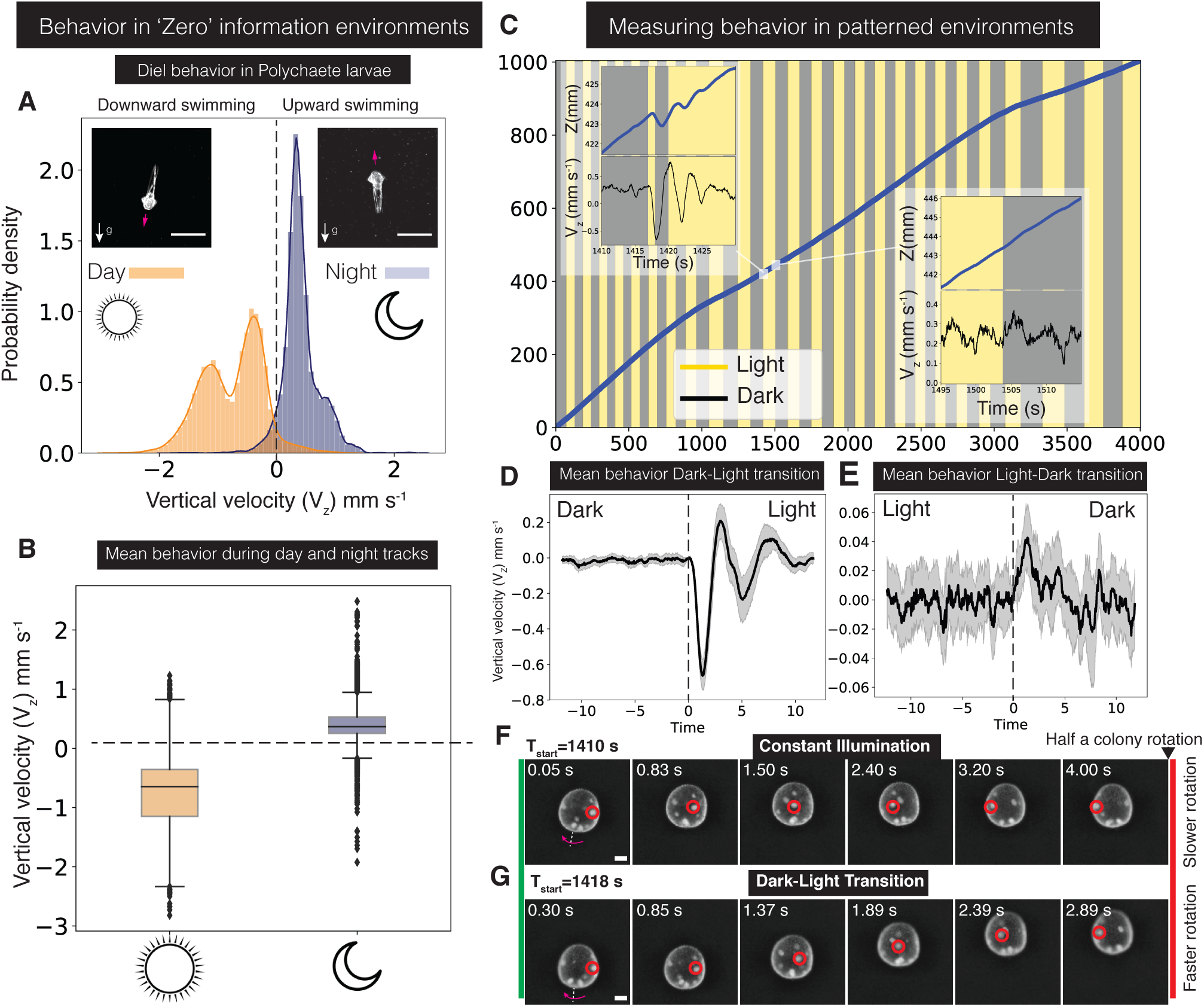
Diel migration at the scale of individual plankton. **(A-B) (A)** Diel swimming behavior of Polychaete larvae (*Owenia sp.*) observed by tracking at day-time (between 1 PM and 3PM local time) and night-time hours (between 10 PM - 11 PM local time). Vertical velocity distributions showed significant differences with swimming velocities being downwards and upwards, at day and night, respectively, with all other experimental conditions maintained constant (Movie 6) (inset, larvae images during day and night tracks, Scale bar, 250 *µm*). **(B)** Vertical velocities over several tracks of different organisms (*n* = 6 organisms, day time, total track duration 360 *s* and *n* = 5 organisms, night time, total track duration 617 *s*). **Behavior in depth-patterned environments**. **(C-G)** Swimming behavior of the colonial algae *Volvox aureus* with ambient light conditions patterned, such that a change in ‘virtual-depth’ of 20 *mm* triggers a Dark-to-Light (D-L) or Light-to-Dark (L-transition. **(C)** A track of a single Volvox colony swimming ≈ 1*m* showing 25 transitions each of D-L and L-D. Microscale behvioral changes accompany the D-L transitions (Left inset, vertical displacement and vertical velocity) whereas the swimming behavior remains unchanged during a L-D transition (Right inset, Movie 7). **(D)** Behavior averaged across 25 D-L transitions by aligning vertical velocity time-traces at the first D-L transition point (mean, solid black line and 95% confidence interval, filled gray region) shows the highly repeatable nature of these transitions. **(E)** In contrast, the behavior at a L-D transition is relatively unaffected. **(F)** Under constant illumination (Light or Dark) *V. aureus* colonies rotate about their swimming direction at a constant rate of about 0.125±0.014*s*^−1^, measured by tracking the daughter cells inside the colonies (red circles). **(G)** During a D-L transition the vertical velocity change is accompanied by a transient increase in the colony rotation rate to 0.217 ± 0.016 *s*^−1^. Scale bars, 100 *µm*.

## Measuring behavior in depth-patterned environments

Organisms migrating in a stratified ocean encounter changes in physical parameters, such as light. By mapping variations of ambient parameters to temporal variations and/or variations in ‘virtual-depth’ of the tracked organism, we can program arbitrary depth-patterned environments. This technique enables us to program both extremely sharp but also extremely shallow (ecological-scale) gradients. We demonstrate this environmental patterning using a light cue that is coupled to a virtual-depth parameter using an ambient light controller (see Fig. 1D). For the purposes of demonstration, we chose an artificial, depth-based light-intensity pattern where every change in depth of 20 *mm* triggered alternately a light-dark (L-D) or dark-light (D-L) transition in the ambient white illumination light intensity, resulting in a “virtual reality arena” such that the organism’s motions directly control its ambient environment. As test subjects, we used the colonial alga *Volvox*, which are made up of hundreds of somatic ciliated cells, and which have a well documented phototactic response [63].

We tracked individual colonies of *Volvox auereus* as they navigated a depth-patterned light environment and triggered alternating D-L and L-D transitions as a function of virtual-depth (Fig. 6C). By measuring individual colonies over long times we were able to measure a number of these transitions (*n* = 25) as the colony swam upwards by 1 *m*. Though not visible over the vast vertical scale of the track, our concurrent microscale measurements revealed that the transition from D-L had a significant effect on the swimming behavior causing the colony to transition from upward swimming to a brief sinking period, which resulted in a second series of L-D and D-L as the colony sank below the depth-based threshold for the transition (Fig. 6C, Top inset; Movie 7). In contrast, the L-D transition did not cause any significant change in the swimming behavior (Fig. 6C, Bottom inset; Movie 7). This asymmetric response towards D-L and L-D transitions was confirmed by measuring the mean behavior, by aligning the vertical velocity traces of several transition events, which displayed universal profiles for D-L and L-D transitions (Fig. 6D, E). The measured mean behavior also confirmed that the brief sinking period triggered by the D-L transition was short-lived and the colony resumed its upward swimming, which is consistent with a transient change in ciliary beat frequency, and a subsequent adaptive response of the cilia, as reported by others using different methods [63]. The transition in the ciliary beat form had a second signature, not reported previously, whereupon the natural rotation frequency of the colonies was perturbed. During a D-L transition the colony transiently rotated faster with *ω_transition_* = 0.217 ± 0.016*s*^−1^ (*n* = 10 transitions) compared to the resting rotation rate during either constant-light or dark conditions *ω_resting_* = 0.125 ± 0.014*s*^−1^ (see Movie 7).

## Single cell tracking

The planktonic organisms considered thus far owe their remarkable motility to cilia that propel them through the fluid. However, a large class of ecologically important planktonic single-celled organisms, such as diatoms and several dinoflagellates, lack any motile appendages. How such cells, being heavier than ambient sea water, do not just sink to the bottom of the ocean is an intriguing problem in cell physiology and ecology. Since these organisms rely on photosynthesis, depth regulation is an important constraint. As a final demonstration of our tracking method, we present two vignettes, one each for diatoms and dinoflagellates, where we connect single-cell behavior and physiology to its mesoscale consequences.

We tracked individual centric diatom cells (Genus Coscinodiscus) over long times (45 *min*), and found a highly dynamic sinking behavior consisting of slow and fast-sinking phases (Fig. 7A,B; Movie 8). The cells are found to transition rapidly (𝒪(10 *ms*) time scales) between these two sinking phases while spending much longer in each phase (fast sinking phase duration: 28.9 ± 28.5 *s*, slow sinking phase: 109.6 ± 65.0 *s*). The distribution of sinking speeds is bi-modal, pointing to an underlying two-state system that controls the cellular buoyancy (Fig. 7C). Concurrent measurements of the flow field around sinking cells (Methods) revealed a conserved hydrodynamic signature during both slow and fast sinking phases (Fig. 7D). This hydrodynamic signature was that of a point force acting downward on a fluid due to the excess density of the suspended cells compared to the ambient fluid. The flow signature during faster sinking corresponded to that of a stronger point-force compared to the slower sinking phase, pointing to a change in cell density (estimated mean of excess cell density Δ*ρ_cell|fast_* = 12.4 *kgm*^−3^ and Δ*ρ_cell|slow_* = 0.8 *kgm*^−3^), as the underlying cause for the change in sinking speeds, even though the exact mechanism is still unknown [64].

**Figure 7:**
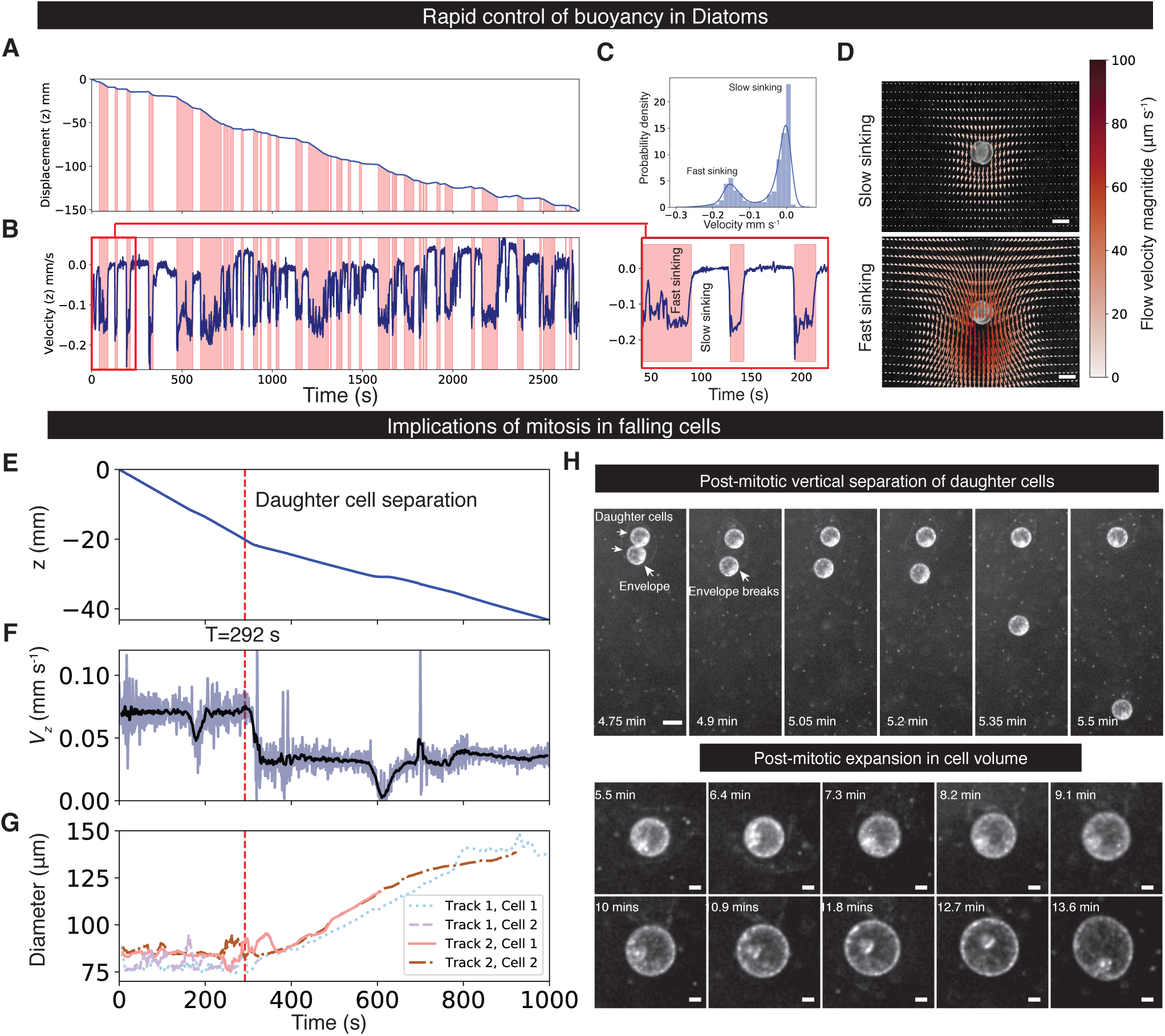
Tracking single cells. **Active control of buoyancy in diatoms (A, B)** Time traces of vertical displacement and vertical velocity of a single, freely sinking diatom cell over long times (45 *mins*), showing a highly dynamic sinking behavior (see Movie 8), consisting of slow and fast sinking phases each lasting 109.6 ± 65.0 *s* and 28.9 ± 28.5 *s*, respectively. **(C)** Sinking speeds show a bi-modal distribution with peaks at the slow and fast sinking-speeds. High-resolution measurements reveal an asymmetry in the transition between the fast to slow sinking which is more gradual than the rapid transition from slow to fast sinking. **(D)** Flow fields around single diatom cells during the slow and fast sinking phases show a similar hydrodynamic signature, which is that of a point force acting on fluid due to excess cellular density, with a larger force for the fast sinking phase, pointing to a change in cell density as the underlying cause of the dynamic sinking behavior. Scale bars, 100 *µm*. **Mesoscale implications of the cell cycle in the dinoflagellate *P. noctiluca***. **(E) and (F)** Time traces of vertical displacement and velocity of a *P. noctiluca* cell completing cell division. The tracked individual daughter cell post-division (vertical dashed line) sinks at a slower rate than before separation. **(G) and (I)** Concurrently the daughter cells undergo a dramatic expansion to roughly twice their initial diameter over a time scale of 10 *mins* (Movie 9). Scale bars, 20 *µm*. **(H)** An asymmetric behavior is also observed in the daughter cells post-division, as one cell remains attached to the ruptured membrane. This asymmetry results in a sinking rate differential between the two daughter cells causing them to become vertically separated. Scale bar, 100 *µm*.

Organisms that spend their life as plankton suspended in the ocean, also undergo basic biological processes such as cell division without ever encountering a solid substrate. It is interesting from the perspective of both cell biology and ecology to understand the changes in sinking rates due to physiological transitions that occur during cell division. To study this we tracked single cells of the dinoflagellate *Pyrocystis noctiluca* as they underwent mitosis while sinking freely. We found that the division process was characterized by a phase where the daughter-cells remained together within a cellular envelope even after cytokinesis (Fig. 7H; Fig. S14). This phase was followed by a rupture of this cellular envelope and ejection of one of the daughter cells, while the other remained attached to the envelope (Fig. 7H). Interestingly, the sinking rate decreased from a higher value of this daughter-cell-envelope complex to a lower one for the individual daughter cells after envelope rupture (Fig. 7E and F). Remarkably, we discovered that post separation, the daughter cells synchronously increased their diameter to about twice that of pre-separation (*d_original_* ≈ 75*µm* to *d_final_* ≈ 140*µm*, a ≈ 6.5*×* increase in cell volume), with a surprising lack of change in the sinking speed (Fig. 7G and I). The daughter-cell separation also showed a pronounced asymmetry where one daughter cell was ejected out of the cellular envelope while the other cell remained in contact with the envelope (*n* = 5 tracked divisions). This asymmetry resulted in a differential in sinking speed, which persisted for as long as one of the daughter cell remained attached to the cell envelope, leading to a significant vertical separation post division (Movie 9). In contrast, conventional imaging of cell-division did not offer any insight into this meso-scale separation along the vertical direction, or density variations during the division process (Fig. S14, Movie 9). This cell-scale process offers a novel mechanism for clonal populations of cells to disperse, with possible ecological consequences. Such measurements of single cells allow us to relate basic cell-scale physiological processes to their effects on buoyancy and hence depth regulation of single-celled plankton.

## Discussion

Conventional microscopy platforms are typically geared towards adherent cells or organisms on cover-slips. However, a large fraction of marine microscale plankton, and ecologically relevant marine particles, spend most of their life-time far from such solid substrates, suspended in the ocean. Furthermore, key processes in the ocean crucially involve free movement and behavior of these discrete entities, in the context of a vertically stratified environment. Imaging such entities under a cover-slip in the horizontal plane, therefore, highly restricts active behavior along the vertical axis, and results in strong interactions with boundaries. This, at best, results in significant modifications to natural behavior, and at worst, leads to important phenomena being completely lost.

The vertical tracking microscopy method presented here, allows one to transcend scales between microscale physiology and ecology by using high-resolution microscopy and allowing free movement along the vertical axis. This method not only allows mapping of complex behavioral trajectories of suspended cells and animals, but also enables simultaneous quantitative recording of microscale properties and physiological state. For instance, density of suspended organic and inorganic particles, as well as organisms is a crucial parameter in understanding vertical material fluxes in the ocean. However, bulk measurements of density don’t offer the full picture since, in many cases, there are several biophysical mechanisms operating at many different scales that affect this density, as demonstrated by our measurements of sinking marine detritus, diatoms and dinoflagellates. We have demonstrated here a means to precisely measure density variations at the scale of individual particles or organisms over ecological scales.

Similarly, by expanding the optical capabilities of the tracking microscope, by adding modules for epi-fluorescence, light sheet, phase contrast or x-ray microscopy, we foresee measurements that make a direct link between planktonic cell’s physiological state, such as the phase of cell-cycle, and its depth in the water column. In plankton that possess a nervous system, we foresee that our method will allow optical recording of individual neurons, while concurrently measuring behavior, which has been demonstrated previously using horizontal tracking methods for other organisms [20, 23, 65].

More broadly, a large space exists for extending the capabilities of the microscope and methodology presented here. In particular, certain optical systems including fluorescence and Differential Interference Contrast (DIC) capability can be readily added, while other systems can be added with changes to the microscope’s design. For such implementations, it may be more suitable to have all motion compensation applied on the circular fluidic chamber so that the optical system may be fixed in the lab frame.

To tease apart the inner mechanisms of how multiple environmental cues (such as light, pressure, salinity etc.) are processed in a plankton, virtual reality experiments with contradictory or correlative environmental landscapes can be constructed. While we have already demonstrated this for the case of varying light intensity as a function of depth, extending this method to include light spectrum, polarization and direction are also readily possible. In the future, we also envision patterning other environmental cues such as chemical concentration and hydrostatic pressure which can be achieved by maintaining fluidic access to the rotating fluidic chamber [29]. Techniques presented here enable an entirely new class of plankton behavioral experiments not possible before.

While we have developed a system geared towards measuring microscale objects, a similar approach of using a “hydrodynamic treadmill” can be used to study macroscale plankton. In an interesting convergence of ideas, while preparing this manuscript, we found an earlier effort where a circular chamber was used, along with manual observation and input by a human operator, to track *macroscale* plankton [66]. In the future, such earlier efforts could be extended by including modern electronics and imaging systems to create an autonomous vertical system for macroscale plankton.

While our method allows scale-free tracking along the vertical direction, the presence of walls in the two horizontal directions, at present, seems unavoidable. Therefore, this method is particularly suitable for objects with anisotropic motility where the dominant motion is along the vertical and is unsuitable for objects with isotropic motility. Gravity, however, ensures that such anisotropy is a rule rather than an exception for earth-bound systems, being especially true for marine microscale plankton [10, 11].

Many fundamental questions remain about key microscale processes in the ocean. How do single-celled plankton sense and regulate their depth? How do plankton with no motile appendages remain suspended in the water column and not sink into the abyss? What sets the rate at which marine particles sediment? How do physical and biological forces shape this falling ecosystem? What is the behavioral space of planktonic larvae, what are the connections between morphology and behavior? This partial list of questions in biological oceanography highlights the urgent need for new tools to study these inherently multi-scale problems. We anticipate tools and techniques presented here will provide a new window into the secret life of plankton and oceanic ecosystems at the finest resolution by finally freeing plankton from the confines of the cover-slip.

## Acknowledgements

We would like to thank Prof. Chris Lowe and Paul Bump for providing access to marine larvae, and Hopkins Marine Station for access to lab space. We would also like to thank the Puerto Rico Science Trust and Isla Magueyes Marine station for lab access. We would like to thank Elgin Korkmazhan for technical assistance for an early version of the experiment. We would like to thank Rebecca Konte for making the drawings of marine larvae and organisms. We thank Prakash lab members for fruitful discussions. D.K was funded by a Bio-X Bowes Fellowship, H.L. was funded by a Bio-X SIGF Fellowship. M.P. acknowledges support from NSF Career Award, Moore Foundation, HHMI Faculty Fellows Program, NSF CCC program and CZ BioHub Investigators program. Authors contributions: D.K. and M.P. designed the research. D.K., H.L., F.B., A.L. and M.P. designed the instrument. D.K., H.L., F.B., P.C. and A.L. built the setup. D.K., H.L., F.B., P.C., A.L. and M.P. performed experiments. D.K. and F.B. performed numerical simulations. D.K. performed analytical calculations. D.K., H.L. and F.B. wrote the control software. D.K., A.L., and M.P. analyzed the data. D.K., H.L., A.L. and M.P. wrote the manuscript. Competing interests: Portions of the technology described here are part of U.S patent-pending (US20190000044A1) to D.K and M.P. Data and materials availability: All data is available in the manuscript or the supplementary materials, and is also available upon request.

## List of Supplementary Materials

Materials and Methods

Supplementary Text

Figs. S1 - S14

Tables. S1 - S3

Movies. S1-S9

## References

[1] R. G. Williams, M. J. Follows, Ocean Dynamics and the Carbon Cycle (Cambridge University Press, 2012).

[2] Y. M. Bar-On, R. Phillips, R. Milo, Proceedings of the National Academy of Sciences 115, 6506 (2018).

[3] P. Field, C. B., Behrenfeld, M. J., Randerson, J. T. and Falkowski, Science 281, 237 (1998).

[4] S. Menden-Deuer, T. Kiørboe, Journal of Plankton Research 38, 1036 (2016).

[5] E. Haeckel, Art forms in nature (Courier Corporation, 2012).

[6] F. Azam, Science 280, 694 LP (1998).

[7] H. L. Bourne, J. K. B. Bishop, T. J. Wood, T. J. Loew, Y. Liu, Biogeosciences Discussions pp. 1–27 (2018).

[8] E. Fuchs, J. Jaffe, R. Long, F. Azam, Optics Express 10, 145 (2012).

[9] D. Yoerger, et al., American Geophysical Union, Ocean Sciences Meeting 2016, abstract #IS14A-2298 (2016).

[10] R. Schuech, S. Menden-Deuer, Limnology and Oceanography: Fluids and Environments 4, 1 (2014).

[11] M. A. McManus, C. B. Woodson, Journal of Experimental Biology 215, 1008 (2012).

[12] D. Blasco, Marine Biology 46, 41 (1978).

[13] G. C. Hays, Migrations and Dispersal of Marine Organisms, M. B. Jones, et al., eds. (Springer Netherlands, Dordrecht, 2003), pp. 163–170.

[14] A. L. Alldredge, M. W. Silver, Progress in Oceanography 20, 41 (1988).

[15] A. L. Shanks, J. D. Trent, Deep Sea Research Part A, Oceanographic Research Papers 27, 137 (1980).

[16] R. K. Cowen, S. Sponaugle, Annual Review of Marine Science 1, 443 (2008).

[17] F. Gross, E. Zeuthen pp. 382–389 (1948).

[18] D. von Wangenheim, et al., eLife 6 (2017).

[19] H. C. Berg, Review of Scientific Instruments 42, 868 (1971).

[20] D. H. Kim, et al., Nature Methods 14, 1107 (2017).

[21] T. Darnige, N. Figueroa-Morales, P. Bohec, A. Lindner, E. Clément, Review of Scientific Instruments 88 (2017).

[22] K. Drescher, R. E. Goldstein, N. Michel, M. Polin, I. Tuval, Physical Review Letters 105, 1 (2010).

[23] L. Cong, et al., eLife 6, 1 (2017).

[24] H. Ploug, B. B. Jørgensen, Mar. Ecol. Prog. Ser. 176, 279 (1999).

[25] T. Sauma-Pérez, C. G. Johnson, L. Yang, T. Mullin, Journal of Fluid Mechanics 847, 119 (2018).

[26] E. A. Van Nierop, et al., Journal of Fluid Mechanics 571, 439 (2007).

[27] K. Mukundakrishnan, H. H. Hu, P. S. Ayyaswamy, Journal of Fluid Mechanics 599, 169 (2008).

[28] A. Wolf, R. P. Schwarz (1999).

[29] R. P. Schwarz, T. J. Goodwin, D. A. Wolf, Journal of Tissue Culture Methods 14, 51 (1992).

[30] S. N. Fry, N. Rohrseitz, A. D. Straw, M. H. Dickinson, Journal of Neuroscience Methods 171, 110 (2008).

[31] K. Drescher, K. C. Leptos, R. E. Goldstein, Review of Scientific Instruments 80, 1 (2009).

[32] C. Verasztó, et al., eLife 7, 1 (2018).

[33] E. A. Salzen, Experimental Cell Research 12, 615 (1957).

[34] L. G. Leal, Advanced transport phenomena: fluid mechanics and convective transport processes, vol. 7 (Cambridge University Press, 2007).

[35] T. Pedley, Annual Review of Fluid Mechanics 24, 313 (1992).

[36] T. J. Pedley, J. O. Kessler, Proceedings of the Royal Society B: Biological Sciences 231, 47 (1987).

[37] R. Hemmersbach, D. Volkmann, D. P. Häder, Journal of Plant Physiology 154, 1 (1999).

[38] A. M. Roberts, Journal of Experimental Biology 53, 687 (1970).

[39] H. C. Berg, Random walks in biology (Princeton University Press, 1993).

[40] J. O. Kessler, Nature 313, 218 (1985).

[41] O. X. Cordero, R. Stocker, Environmental Microbiology Reports 9, 16 (2017).

[42] P. S. Ayyaswamy, K. Mukundakrishnan, Acta Astronautica 60, 397 (2007).

[43] E. Santamaría, B. Ayo, J. Iriberri, A. López, I. Artolozaga, Journal of Plankton Research 19, 1429 (2007).

[44] G. Herndl, Marine Ecology Progress Series 48, 265 (2007).

[45] H. Ploug, H. P. Grossart, F. Azam, B. B. Jørgensen, Marine Ecology Progress Series 179, 1 (1999).

[46] A. F. Fortes, D. D. Joseph, T. S. Lundgren, Journal of Fluid Mechanics 177, 467 (1987).

[47] M. Cable, J. R. Frade, Journal of Materials Science 22, 1894 (1987).

[48] M. Cable, D. J. Evans, Journal of Applied Physics 38, 2899 (1967).

[49] N. Shankar, T. J. Wiltshire, R. Shankar Subramanian, Chemical Engineering Communications 27, 263 (1984).

[50] R. R. Strathmann, Integrative and Comparative Biology 15, 717 (1975).

[51] R. R. Strathmann, D. Grünbaum, Integrative and Comparative Biology 46, 312 (2006).

[52] T. C. Lacalli, T. H. J. Gilmour, J. E. West 330, 371 (1928).

[53] J. T. Pennington, R. R. Strathmann, Biological Bulletin 179, 121 (1990).

[54] R. B. Emlet, Marine Ecology Progress Series 63, 211 (1990).

[55] P. Gonzalez, J. Z. Jiang, C. J. Lowe, Frontiers in Zoology 15, 1 (2018).

[56] G. O. Mackie, A. N. Spencer, R. Strathmann, Nature 223, 1384 (1969).

[57] T. Fenchel, K. W. Ockelmann, Ophelia 56, 171 (2002).

[58] W. Gilpin, V. N. Prakash, M. Prakash, Nature Physics 13, 380 (2017).

[59] B. Pernet, Evolutionary Ecology of Marine Invertebrate Larvae (2018), vol. 1.

[60] M. Conzelmann, et al., Proceedings of the National Academy of Sciences 108, E1174 (2011).

[61] G. Jékely, et al., Nature 456, 395 (2008).

[62] M. A. Tosches, D. Bucher, P. Vopalensky, D. Arendt, Cell 159, 46 (2014).

[63] K. Drescher, R. E. Goldstein, I. Tuval, Proceedings of the National Academy of Sciences 107, 11171 (2010).

[64] B. J. Gemmell, G. Oh, E. J. Buskey, T. A. Villareal, Proceedings of the Royal Society B: Biological Sciences 283 (2016).

[65] J. P. Nguyen, et al., Proceedings of the National Academy of Sciences (2015).

[66] A. C. Hardy, R. Bainbridge, Journal of the Marine Biological Association of the United Kingdom 33, 409 (1954).

