## Supplementary Material for "Scale-free Vertical Tracking Microscopy: Towards Bridging Scales in Biological Oceanography"

April 15, 2019

**3 Supplementary Movie Captions** **34**

### 1 Materials and Methods

#### 1.1 Circular Fluidic Chamber Construction

The Circular fluidic chamber was custom fabricated from four parts: two spacer rings machined out of aluminum which were sandwiched between clear, scratch-resistant acrylic sheets, resulting in an annular volume which was optically accessible from both sides (Fig. S1B, C). The dimensions of the resulting annular volume was given by its inner radius $R_i$, outer radius $R_o$, and width $W$. Typical values of these parameters used in our experiments were $R_i = 85\ mm$, $R_o = 100$ to $115\ mm$ and $W = 3$ to $6\ mm$ ( Fig. S1D), resulting in cross-sectional dimensions $L \times W$, where $L = R_o - R_i$. 100 % silicone adhesive (GE silicone) was used to bond the various layers of the AFC so as to provide a biocompatible seal which is also gas permeable to allow long-term experiments. Inlet and outlet ports made via luer attachments (Cole-Parmer) allowed the chamber to be completed filled with fluid and also allowed objects of interest to be introduced.

#### 1.2 Motorized stages for tracking

The circular fluidic chamber was attached to a fine rotational stage using a high precision shaft (Phidgets Inc.) and rotational bearings (Robotshop Inc., Canada) via a torsional-beam coupler (Pololu robotics) (Fig. S1 A). The rotational stage comprised of a NEMA-11 stepper motor mated to a 100:1 gearbox (Phidgets Inc., Canada), with a horizontal rotational axis (Fig. S1 A). This arrangement allowed the chamber to be rotated with an angular resolution of $19 \pm 1\ \mu\ radians$ per step, resulting in a tangential linear increment of $1.73 \pm 0.08\ \mu m$ per step at the center-line of the annulus. The rotation of the stage was measured using optical encoders (Phidgets Inc.) with a resolution of $105.2\ \mu\ radians$ per pulse. The rotational stage and bearings were mounted on height adjustable posts (ThorLabs, New Jersey, USA) which were adjusted to ensure a horizontal axis.

The two other motion axes ($x$ and $y$) were implemented using either standard off-theshelf translational stages driven by stepper motors (Haijie Technology Ltd., Beijing). In our current implementation, the optical assembly was attached to the $xy$ stages, such that tracking in the $xy$ directions was achieved by translating the optical assembly to follow the object. Alternative designs are possible where motion compensation along all 3 axes (2 translation and 1 rotation) are applied to the circular fluidic chamber such that the optical system can be fixed in the lab reference frame. Such an implementation may be better suited when we one is interested in building advanced microscopy systems to work in conjunction with our tracking method.

#### 1.3 Optical system

We constructed a light microscope focused on either the 3 O' clock or 9 O' clock position of the circular fluidic chamber such that rotational motion of the chamber resulted in approximately vertical motion (along the $z$ direction) in the optical field-of-view (Fig. S1A). The optical assembly was mounted on motorized translational $xy$ stages for motion compensation in the horizontal directions. The optical assembly consisted of a lens assembly with an incorporated liquid-lens (Corning, Varioptic) which served as the imaging objective (finite conjugate configuration), coupled to a CMOS camera (DFK 37BUX273, The Imaging Source, Germany), capable of full resolution imaging (1440x1080 pixels) at 238 Hz, resulting in an optical FOV of $2293\mu m \times 1720\mu m$. The imaging system was modular so that different modalities could be interchanged. For tracking we used dark-field (DF) imaging using a ring LED assembly situated on the opposite side of the fluidic chamber (Fig. S1). We primarily used red (625 $nm$) (Fig. S12) LEDs to image since this does not induce phototactic behaviors in most organisms. Images captured on the camera sensor were processed using a custom image-processing pipeline implemented on a standard desktop CPU at rates of 100 Hz.

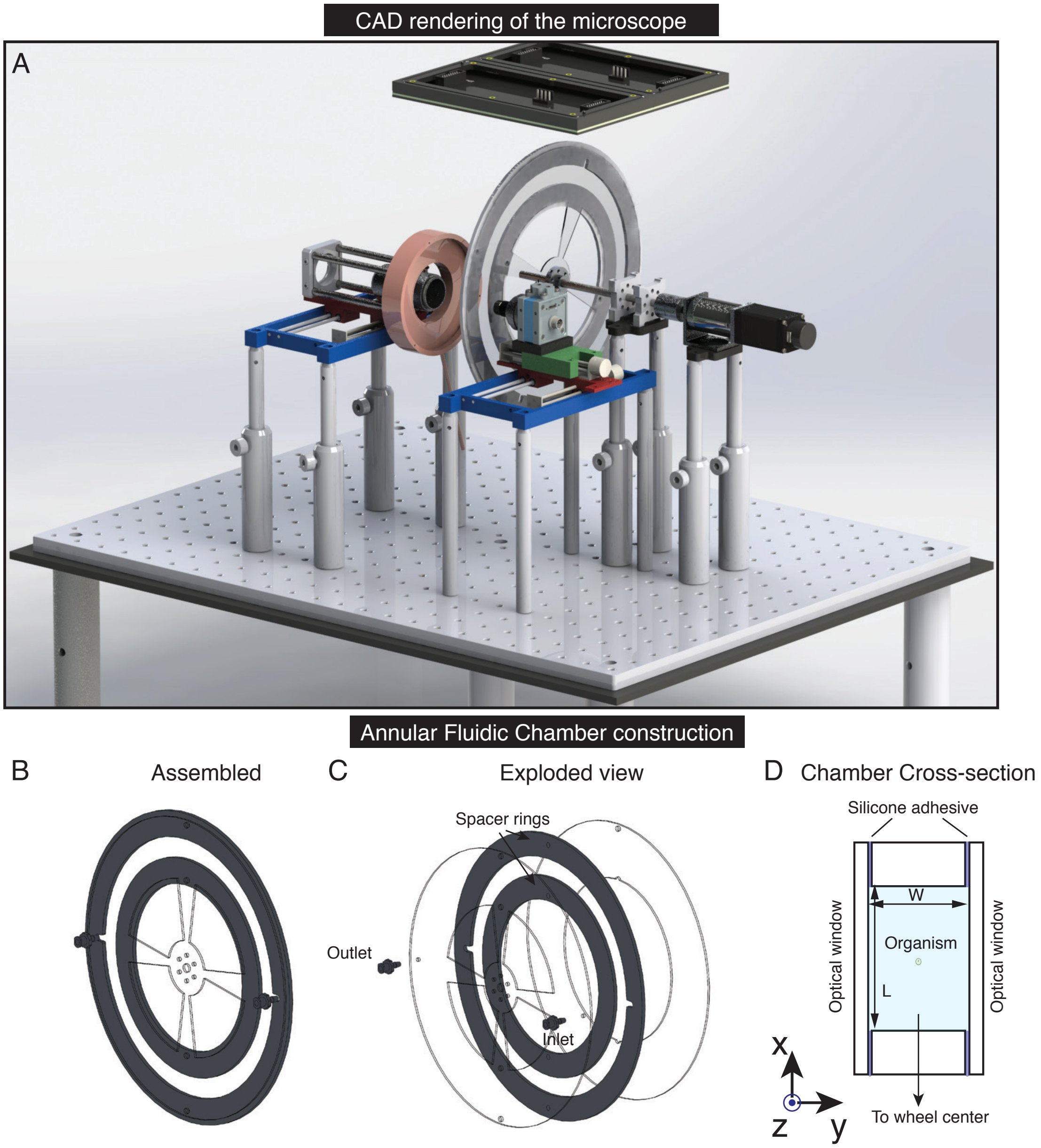

**Figure S1** **(A)** CAD rendering of the tracking microscope, showing all major components as well as ambient light controller. **(B)** Assembled and **(C)** exploded view of the circular fluidic chamber. **(D)** Cross-section of the circular fluidic chamber.

#### 1.4 Control system for tracking

Images for tracking were obtained from the CMOS sensor at rates of 100 Hz, and processed using a custom image processing pipeline implemented using using Python on a desktop CPU (Fig. S2A). This pipeline consisted of separate organism trackers for lateral ($xz$) and axial ($y$) positions. For lateral positions, an initial region of interest containing the object was selected by the user, and this object was tracked in further frames using an open-source object tracking algorithm available on OpenCV-Python [1] (Fig. S3). This algorithm was robust enough to track the same object over a long times (1 day), even in the presence of other similar looking objects, organisms and debris. As an alternative, to take advantage of computers that have a GPU (Graphics Processing Unit), we utilized a hardware accelerated object tracking algorithm [2] which allowed a combination of robust tracking and high frame rates (up to 200 Hz). In all cases the output of the organism tracker was the lateral position $(x_{obj}, z_{obj})$ of the object relative to the center of the microscope's FOV. This output was fed through a PID controller which in turn calculated the error signals that were sent to the motorized stages (Fig. S2A).

##### 1.4.1 Focus tracking

For estimating the axial position of objects, a separate focus tracking algorithm was developed. This used a liquid-lens (Caspian u-25H0-075, Varioptic, Corning) to rapidly scan the focal plane and obtain image stacks at up to 30 volumes per second over a depth range of 50-500 $\mu m$ (Fig. S2B, C; Fig. S3). A focus measure of an image was estimated using the image intensity variance [3, 4] (Fig. S3). A peak-finding algorithm was further used to determine the focal plane position corresponding to the best focus and hence the object's position. This estimated position was fed to a proportional controller, with a tunable gain and the resulting error signal was used to move the $y$-axis stage to follow the object (Fig. S2C). We characterized the tracking performance of this method by tracking a 250$\mu m$ bead mounted to a motorized stage that allowed prescribed motion of the bead along the

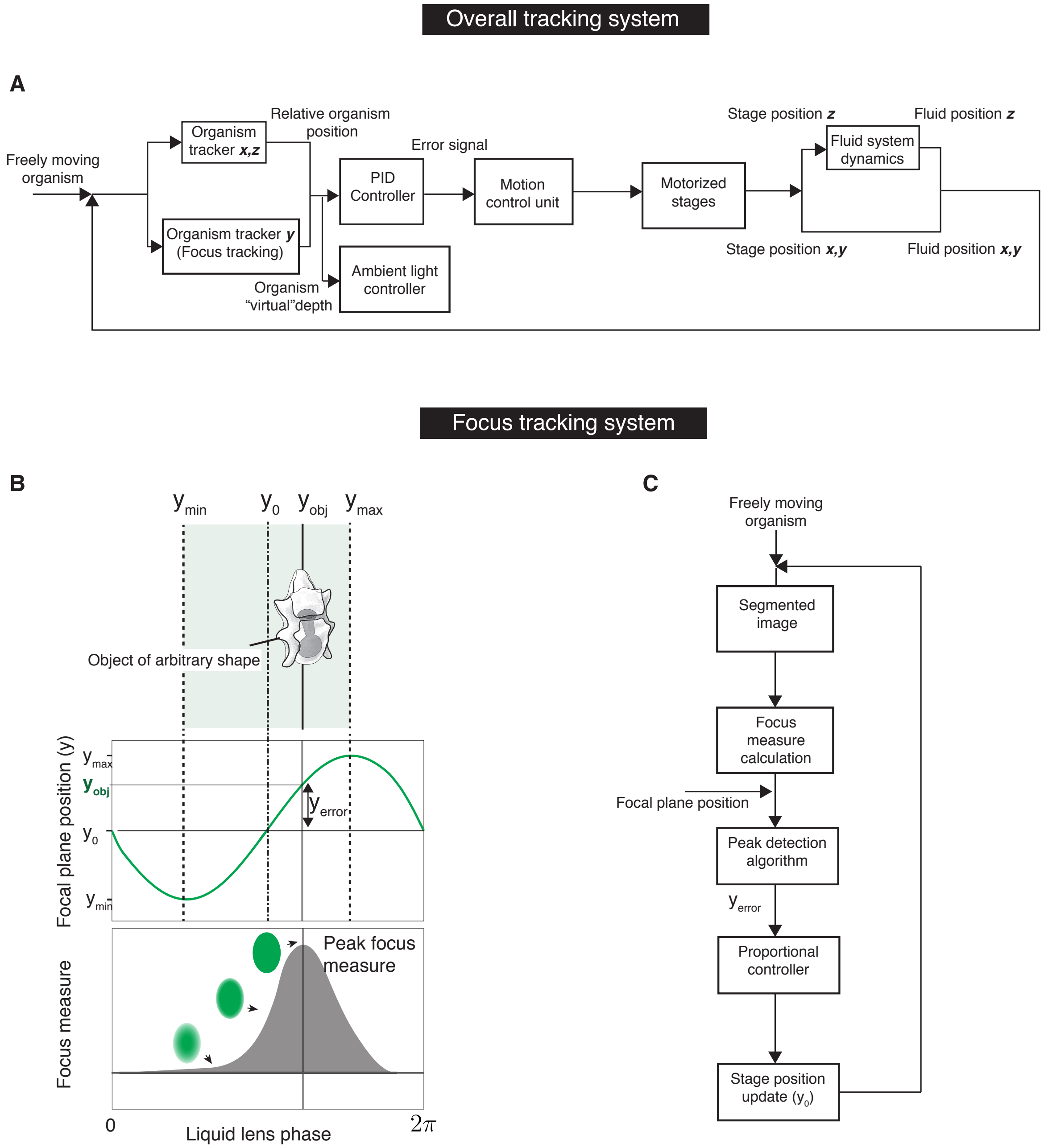

**Figure S2** **(A)** Flow-chart of the tracking control system. **(B)** Focus-tracking methodology using a liquid-lens to sweep the focal plane of the microscope (Top, Middle), while calculating the focus measure (image variance) to locate the optimal focal plane that maximizes this focus measure (Bottom). **(C)** Flow-chart of the focus-tracking sub-system.

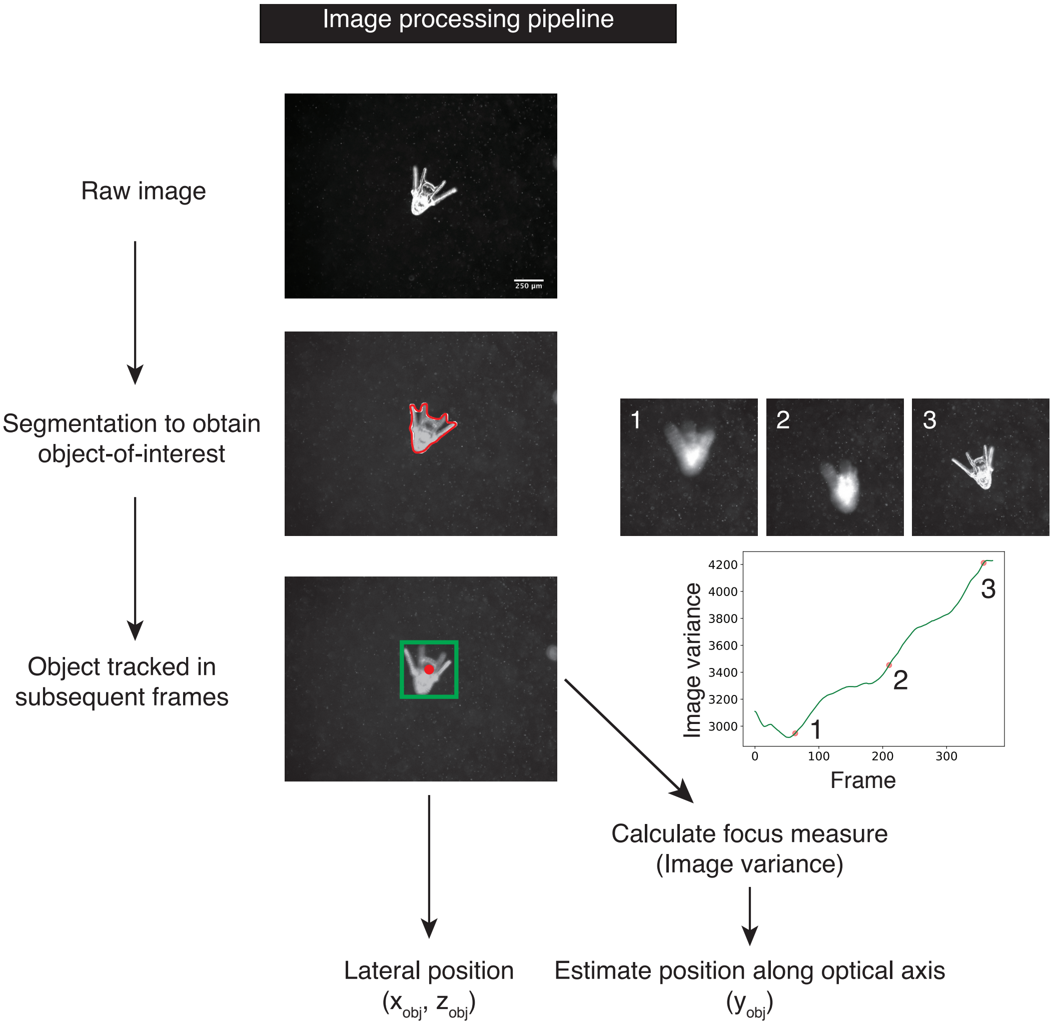

**Figure S3** Image processing pipeline to initially obtain the object's centroid and track it in further frames. The initial location of the object is obtained by using a color-based threshold in HSV-color-space. In further frames the object is tracked using OpenCV's CSRT tracking algorithm [1], or the DaSiamRN algorithm [2]. The sample images are shown for Sand Dollar larvae (*D. excentricus*), which has a complex shape. Focus-measure (image variance) calculation for use in focus-tracking, showing both out-of-focus and in-focus images and their associated focus measures.

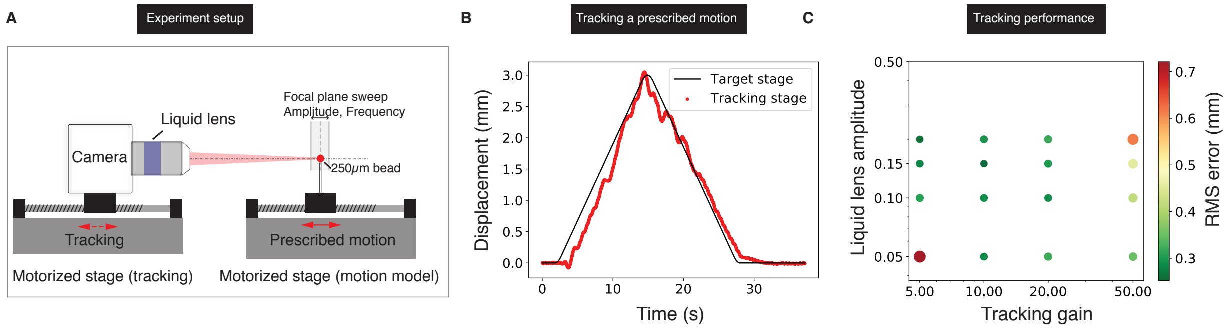

**Figure S4** **(A)** Experimental setup used to characterize the performance of the focus-tracking system. This consists of two motorized stages, the first of which has a $250\ \mu m$ bead attached, and executes a prescribed motion profile. The second stage is the tracking stage with the camera and liquid-lens optical system attached, which tracks the prescribed motion of the bead. (B) Displacement vs time plot for the stage with prescribed motion (black solid line), and the tracking stage (red dots) for liquid-lens amplitude of $100\mu m$ and proportional gain of $10$. **(C)** Characterization of focus-tracking performance showing a plot of the root-mean-square (RMS) error between the tracked and tracking stage as a function of the liquid-lens amplitude and the gain of the proportional controller. By an optimal choice of the lens amplitude and controller gain, RMS errors less than the tracked object's size could be achieved.

optical axis (Fig. S4A). This bead was then tracked using our focus tracking strategy by obtaining volume scans with the liquid-lens and translating the optical assembly to track the object (Fig. S4B). We characterized the tracking performance as a function of the scanning amplitude of the liquid-lens and the tune-able gain, and found an optimal range for these parameters (see Fig. S4C).

The error signals for all three-axes were sent to a Motion Control Unit which was a Arduino-Due microcontroller (Arduino). The microcontroller, in turn, was used to calculate the motion profiles for the motorized stages. These signals were sent to a dedicated stepper motor driver for each motorized stage axis (Big Easy Driver, Sparkfun) which used the Allegro A4988 stepper driver chip. The positions of the stages were measured using rotary optical encoders (HKT22, Phidgets Inc.) with a quadrature resolution of 600 counts per revolution.

#### 1.5 Loading of the circular fluidic chamber and ensuring thermal equilibration

Before experiments the fluidic chamber was passivated by filling it with 5 % BSA solution (Fisher Scientific) and allowing it to sit for 1 hour. After this treatment the fluidic chamber was rinsed twice with the standard solution to be used for the experiments. For the actual experiments, the chamber was completely filled with the appropriate standard solution, through the luer attachments, taking care to avoid the formation of bubbles. Once the chamber was filled, it was mounted on the rotational stage within an enclosure whose temperature was set to $22°C$ using a temperature control unit (AirTherm SMT, World Precision Instruments). During this time, the fluid suspension containing the objects or organisms to be tracked was also stored within this enclosure. After this a fluid mixing protocol was activated to achieve thermal equilibrium between the chamber and fluid. Such a thermal equilibration was necessary in order to prevent thermally driven flows from occurring in the chamber during the experiments. This mixing protocol consisted of a rotational motion of the chamber and periodically reversing the rotation direction. These motions leads to a shear-enhanced mixing of fluid in the chamber, and a correspondingly more effective heat transfer, both between azimuthally separated fluid parcels, as well as between the fluid and the chamber. This mixing protocol was carried out for 10 minutes prior to introducing the objects to be tracked. After this protocol the average background fluid motion when the fluidic chamber was at rest was measured using Particle-Image-Velocimetry (PIV), and was found to typically be $< 20 \mu m s^{-1}$ (Fig. S5). Once thermal equilibrium was achieved, the objects or organisms to be tracked were introduced and the above equilibration procedure was again run for 10 mins to evenly suspend the objects or organisms in the fluid. Once this procedure was completed tracking could be started. This was achieved by manually locating an object of interest and then starting the automated tracker (Movie 1). Care was also taken to ensure significant air-circulation in the experimental enclosure and prevent sources of heat from being present near the fluidic chamber as these can cause thermally driven flows

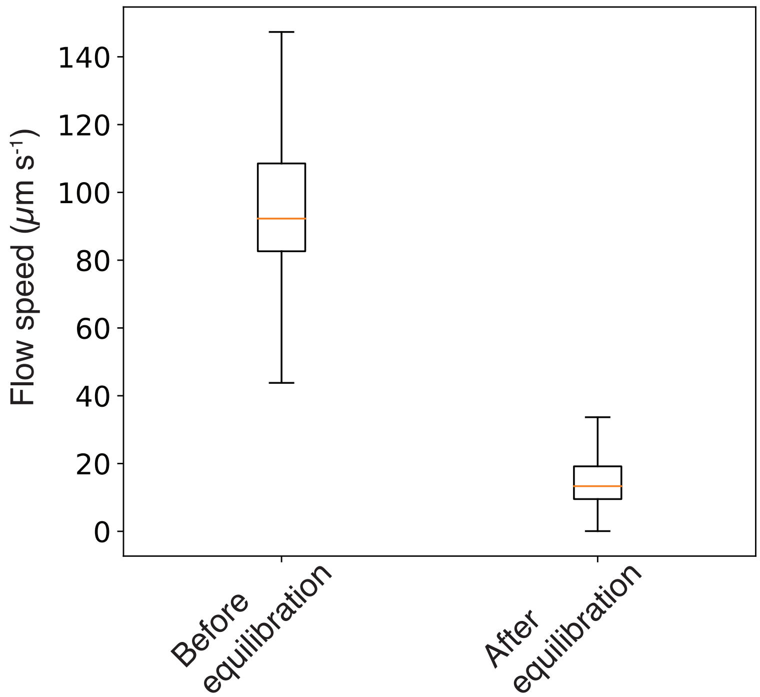

**Figure S5** Background flow measurement. Background flow speeds in the circular fluidic chamber before and after the thermal equilibration procedure. This flow was measured over the microscope FOV for a duration of 6 seconds at 30 Hz using Particle Image Velocimetry. The box-plot is from the lower and upper quartile of the data, the line represents the median and whiskers show the range. Typical flow speeds were measured to be $< 20\ \mu m s^{-1}$ after the thermal equilibration procedure.

to occur during long-term imaging.

#### 1.6 Abiotic experiments

For validating our experiments we used density marker beads of known size and density purchased from Cospheric (Table S2). These beads have a precisely calibrated density which is slightly higher than water, as well as a mono-disperse size range (Table S2)). We measured the sedimentation velocity of these beads as a means to validate our tracking method and microscope. To perform this calibration, these beads were tracked in two ways. As a control, Eulerian tracks of the beads were obtained by allowing them to sediment down a vertical cuvette with a height of 150 $mm$ and cross-sectional dimensions 15 $mm \times 3$ $mm$. The beads

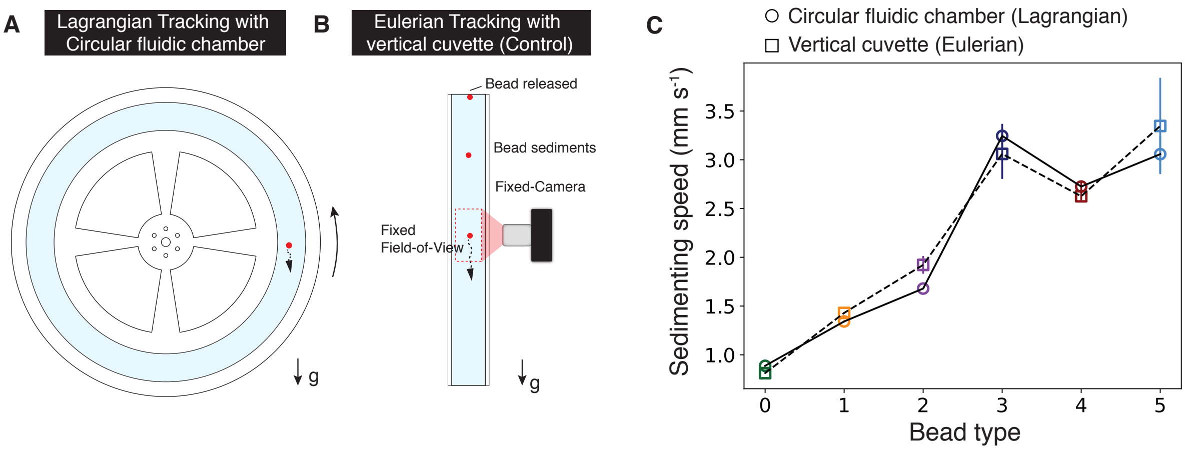

**Figure S6** Validation of the vertical tracking method. We measured the sedimentation speeds of 250 $\mu m$ density calibration beads using **(A)** the method presented in this work and **(B)** conventional fixed-camera method with a vertical cuvette. **(C)** Comparison of measured sedimentation speeds for 6 types of beads with different densities and diameters (data is for $n = 10$ tracks per bead type and tracking method) (see Table S2) using the vertical tracking method presented in this work (circles) and conventional method (squares). Error bars represent one standard deviation.

were imaged as they sedimented past a fixed camera mounted to image in the vertical plane at a location at the middle of the cuvette (see schematic in Fig. S6B), so as to avoid end effects. Lagrangian tracks of the density marker beads were obtained by tracking them using the circular fluidic chamber and tracking methodology developed in this work (Fig. S6A). Each bead was tracked for a 1 $min$ after which the track was stopped and a new bead tracked. The sedimenting velocity, for both the tracking methods above, was obtained by a linear fit of measured time-traces of vertical displacement. The measured sedimentation velocities ($n = 10$ tracks, per bead, per method) are shown in Fig. S6C, and were found to be in good agreement.

The spherical beads used for measuring interactions between sedimenting particles were 500 $\mu m$ solid glass spheres with density 2200 $kgm^{-3}$ (Cospheric LLC., USA). The rods used for the experiments were pencil leads of different lengths in the range $0.5 - 2$ $mm$, diameter 500 $\mu m$ and density (2300 $kgm^{-3}$). The crystals used for the sedimentation-dissolution experiments were raw sugar crystals (Turbinado cane sugar) chosen since they dissolve slower compared to refined sugar crystals of the same size. Marine detritus particles used in experiments were collected during night-time (10 PM local time on 31 August 2018) plankton tows off the coast of Monterey, California, USA using plankton net size of 100 $\mu m$ and isolated under a dissection scope. It was then suspended in filtered sea water from the Hopkins marine station before tracking on the microscope.

#### 1.7 Marine invertebrate larvae experiments

The larvae used for experiments were obtained by fertilizing adult animals collected off the coast of Monterey, California, USA. The culturing procedure for larvae matched standard protocols in the field [5]. Before experiments the larvae were transferred to filtered sea water and allowed to acclimatize for an hour. Larvae were transferred to the circular fluidic chamber using tubing at least three times their body size using gentle suction.

#### 1.8 Environmental patterning experiments with Volvox

*Volvox aureus* colonies were obtained from Carolina Inc. and cultured in Volvox medium (UTEX). For the environmental patterning experiments, we used a white LED array (Adafruit) (Fig. S1A; Fig. S12) mounted at the ceiling of our experimental enclosure, which provided a uniform top illumination at the sample location at the 3 o'clock location of the circular fluidic chamber. The intensity of the LEDs was controlled using a Pulse-Width-Modulation (PWM) signal from the Arduino microcontroller, in turn controlled by the desktop CPU based on the virtual depth of the tracked organism (see Fig. S2A). Using this system any temporal (and hence 'virtual-depth'-based) intensity profile could be programmed into our experiments as a function of the virtual depth of the organism being tracked. In our experiments we simulated an artificial profile which alternated between light and dark for every 20 $mm$ in height gained by the organism. Red (625 $nm$, Fig. S12) LEDs were used to image the organisms since this is a wavelength to which Volvox is insensitive [6].

#### 1.9 Single cell tracking experiments

Pyrocystis noctiluca were obtained from UTEX (UTEX LB 2504) and cultured in F/2 medium (Bigelow Labs) at 20-22°C while maintaining a 12/12H light/dark cycle. Cells were loaded into the chamber 3H after the beginning of the light cycle and allowed to equilibrate for 1H with gentle rotation in the apparatus before tracking. Cells were imaged using red (625$nm$) illumination under a white LED array maintaining the daylight cycle. 3 $\mu m$ latex beads were included for PIV measurements.

#### 1.10 Data analysis procedures

##### 1.10.1 Flow field calculation using Particle Image Velocimetry (PIV)

For calculating flow fields around objects and freely swimming organisms, the fluid was seeded with neutral density tracer particles (2 or 3 $\mu m$ polystyrene beads, Polysciences) with a typical final concentration of 0.05% by volume of beads in the standard solution being used in the particular experiment. For resolving dynamic flow fields a sampling rate of at least 30 Hz was used, and this was adjusted to a higher value based on the flow speeds in different experiments. The images obtained were analyzed using an open-source PIV software implemented on Python [7]. The parameters used for the PIV data analysis are provided in Table S3. The data obtained from the PIV analysis was post-processed as follows. Outlier vectors were detected using the signal-to-noise ratio threshold of the correlation peaks heights, and replaced by the local nearest neighbour average. The data was also interpolated on to a 2x finer grid than that used for calculating the correlations for estimating flows near the surface of organisms. For calculating path-lines of tracer particles in the fluid we used a sliding-window-based maximum intensity projection method as implemented using the Flowtrace software package [8].

#### 1.11 Mapping measured angular displacement of the chamber to virtual depth of object

Our tracking method uses a circular chamber as a 'hydrodynamic treadmill' to track vertical motion over unlimited scale. This method implies that the tracked object does not move relative to the lab reference frame, and its displacement with respect to the fluid is measured by the angular displacement of the chamber. We can map this angular displacement to the 'virtual depth' of the object as follows:

$$Z^{i+1} = Z^i + (Z^i_{com} - Z^{i-1}_{com}) + (R_{center} + X^i)(\theta^i - \theta^{i-1}), \tag{1}$$

where the superscripts $i$ denote discrete time in control-loop cycles, so that $Z^i$ is the 'virtual depth' of the object at time $i$, $Z^i_{com}$ is the centroid of the object relative to the optical Field-of-View (FOV), $R_{center}$ is the radius of the chamber's centerline, $X^i$ is the displacement along the $X$-axis, as measured from the chamber center-line, and $\theta^i$ is the angular displacement of the circular chamber.

The displacement along the $x$ direction is give by:

$$X^i = X^i_{FOV} + X^i_{com}, \tag{2}$$

$$Y^i = Y^i_{FOV}, \tag{3}$$

where $X^i_{FOV}$ is the displacement of the optical FOV relative to the center-line of the chamber and $X^i_{com}$ is the $x$ displacement of the object's centroid compared to the optical FOV. Note that our $y$ displacement is taken as the location of the focal plane of the optical system, which tracks objects movements using our focus tracking system (see §1.4.1). Also note that the FOV along the $x$ and $y$ directions, unlike $z$, are not fixed in the lab reference frame, which is why the update formulae in Eqs. (2) and (3) are different from those in Eq. (1).

### 2 Supplementary Text

#### 2.1 Calculation of relevant non-dimensional numbers for tracking using a circular fluidic chamber

To understand the design space and operating parameters involved in using a circular fluidic chamber as a "hydrodynamic treadmill" to track vertical motion, we performed a scaling analysis of the various physical effects at play. This initial scaling analysis helps us understand which of these physical effects are important, which then motivated us to do a more detailed analysis of those effects on the tracking performance, presented in subsequent sections and also discussed in the main text of this work.

Consider the tracking of objects with a size scale $d$, a vertical speed relative to the ambient fluid of $u_{obj}$, mean density $\rho_{obj}$, in an ambient fluid with density $\rho_f$ and kinematic viscosity $\nu$. In our work, we restrict ourselves to objects or organisms which are small, with sizes which are $\mathcal{O}(mm)$ and smaller. Also these objects have a typical speed relative to the fluid of a few body lengths per second. For the sake of this analysis, we consider a spherical object with diameter $d = 1\ mm$, and speed $u_{obj} = 1\ mms^{-1}$, immersed in water ($\nu \approx 10^{-6}\ m^2\ s^{-1}$. For such an object, the Reynolds number, which quantifies the relative importance of inertial and viscous effects in the flow [9], is:

$$Re = \frac{u_{obj} d}{\nu} \approx 1. \tag{4}$$

This sets an upper bound, at least for the biological organisms considered in this work.

A more relevant dimensionless number for tracking, is the Stokes number which quantifies the relative time scales of the object and the time scale of the flow [9], and is given by :

$$Stk = \frac{t_{obj}}{t_{flow}}, \tag{5}$$

where $t_{obj}$ is the relaxation time-scale of the object, i.e. the time scale over which the object's velocity decays exponentially due to fluid drag, and $t_{flow}$ is a characteristic time-scale of the flow. In our tracking system, the relevant time-scale for the flow is that of viscous diffusion of momentum since this sets the time over which the fluid changes velocity due to changes in the chamber's velocity. This time scale is given by $W^2/\nu$, where $W$ is smallest cross-sectional chamber dimension (the chamber width). The relevant relaxation time-scale for the object, for the case of Stokes flow ($Re < 1$) is given by:

$$t_{obj} = \frac{\rho_{obj} d^2}{18\mu}, \tag{6}$$

where $\mu$ is the dynamic viscosity of the fluid. For most biological organisms their mean density falls in the range of a $5 - 10\%$ excess density to that of water [10]. Assuming these density ranges and the parameters used above for the size and speed, the Stokes number number is $Stk \approx 0.006$. Since $Stk \ll 1$, the object's inertial time-scale is negligible, and the object is advected by the ambient fluid motion. This means that the net motion relative to the fluid is, at leading order, only because of a body forces, such as those due to gravity, and/or active swimming stresses. Smaller, higher order Faxén's corrections exist due to the finite size of the object in a transient non-uniform flow profile [11], which we neglect.

##### 2.1.1 Effects of chamber curvature and rotation

Our next considerations are understanding the role of curvature of the chamber on the tracked object's motion. Firstly, since the tracked object is immersed in a fluid that is globally undergoing a solid-body-rotation, it is subject to an angular velocity given by:

$$\Omega_{rotation} = u_{obj}/R, \tag{7}$$

where $R$ is the object's radial location. Using typical values of $u_{obj} = 1\ mm\ s^{-1}$ and $R = 100\ mm$, we see that $\Omega_{rotation} = 10^{-2} s^{-1}$. We will see in subsequent sections, that this is a negligible contribution to the object's orientation dynamics, and is sub-dominant to the effects of small bottom-heavy density distributions, as well as object shape, both of which control the object's orientations at much faster time-scales.

Chamber rotation gives rise to centrifugal forces which establish a radial pressure gradient, in addition to the hydrostatic gradient due to the vertical orientation of the chamber. The centrifugal contribution to this pressure gradient is given by:

$$\left.\frac{\partial p}{\partial r}\right|_{centrifugal} = \frac{\rho_f u_{obj}^2}{R}, \tag{8}$$

where $R$ is the radial location of the object relative to the center of the chamber.

This radial pressure gradient causes a radial drift of objects embedded in the fluid which can be computed considering the balance of buoyant and fluid drag forces [9]:

$$U_{centrifugal} = \frac{(\rho_s - \rho_f) V_{obj} u_{obj}^2}{C_D \mu d R}, \tag{9}$$

where $V_{obj}$ is the object's volume and $C_D$ is the drag coefficient. For a spherical object, this simplifies to the expression, rewritten as a relative velocity ratio:

$$\frac{U_{centrifugal}}{u_{obj}} = \frac{(\rho_s - \rho_f) d^2 u_{obj}}{18 \mu R}, \tag{10}$$

which for the parameters used above, and $\rho_{obj} = 1100 kgm^{-3}$ is $U_{centrifugal}/u_{obj} = 6.35 \times 10^{-5}$. Thus drift due to centrifugal forces can be safely neglected.

#### 2.2 Hydrodynamic considerations for tracking

Tracking vertical motion using a circular fluidic chamber with a contiguous annulus of fluid (i.e. with no walls normal to the direction of chamber rotation) implies that the fluid has a separate degree-of-freedom and does not rigidly move with the chamber walls. Any change in momentum of the chamber is transmitted to the fluid only via viscous diffusion with an associated temporal delay and spatial non-uniformity in the fluid motion. To achieve effective tracking, we need to account for this fluid motion and understand its effects on

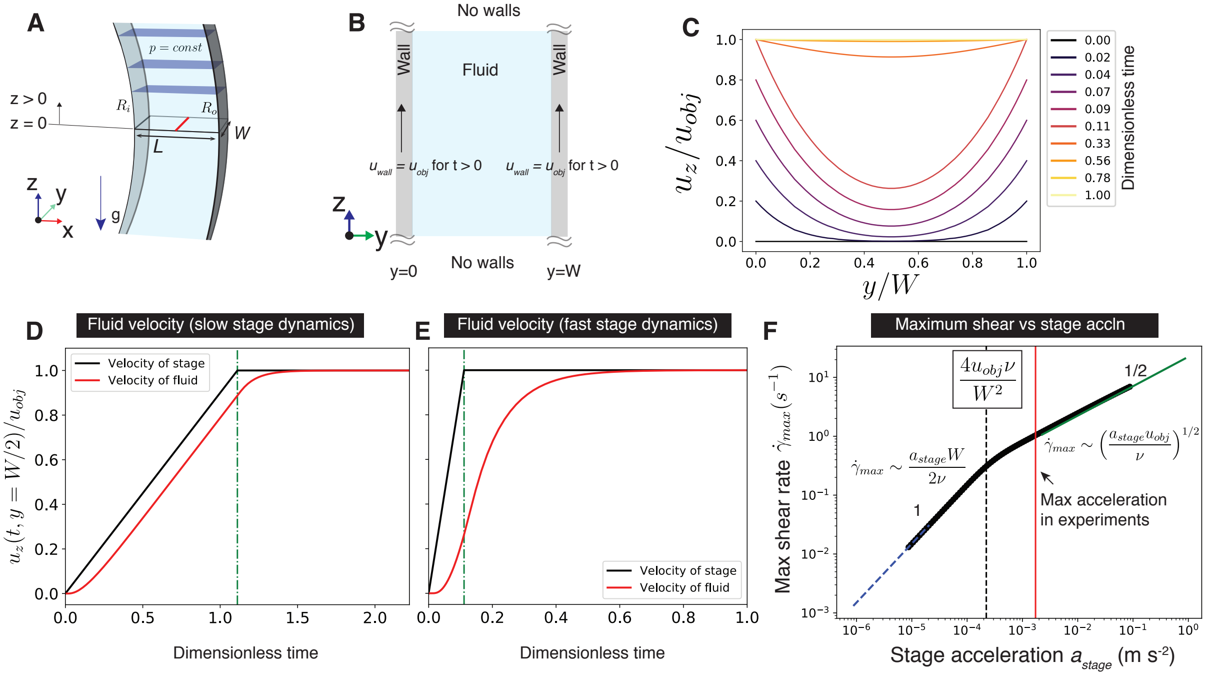

**Figure S7** Hydrodynamic considerations for vertical tracking using a "hydrodynamic treadmill". **(A)** Coordinate system, also showing iso-bars when the fluid is at rest. **(B)** Geometry and boundary conditions for solving for the flow due to impulsive start-up motion of the channel walls. **(C)** Theoretically predicted flow velocity as a function of distance across the channel for an impulsive start-up motion of the channel walls (colored contours: time series of flow velocity profiles). **(D)-(E)** Flow velocity vs dimensionless time ($\tilde{t} = t\nu/W^2$) at the center-line of the channel for the limits of (D) small and (E) large stage accelerations. **(F)** The spatio-temporal maximum in shear rate developed in the fluid during a start-up motion with finite stage acceleration $a_{stage}$. The shear rate has two asymptotic regimes based on the magnitude of the stage acceleration. For slow stage dynamics, compared to the viscous time scale ($\tau_{stage} \gg \tau_{visc}$), the shear rate scales as $\dot{\gamma}_{max} \sim a_{stage}W/(2\nu)$ (blue dashed line), where $\tau_{stage} = u_{obj}/a_{stage}$ and $\tau_{visc} = W^2/\nu$. For fast stage dynamics ($\tau_{stage} \ll \tau_{visc}$), the shear rate scales as $\dot{\gamma}_{max} \sim (a_{stage}u_{obj}/\nu)^{1/2}$ (green solid line). The cross-over between the two scaling regimes is given by solving $\tau_{stage} \sim \tau_{visc}$. The shear-rate corresponding to the maximum stage acceleration (red vertical line) used in our experiments is $\approx 1s^{-1}$.

the tracks obtained. Towards this, we model the dynamics of fluid motion in response to arbitrary control inputs to the chamber.

The fluidic chamber is an annulus with inner and outer radii $R_i$ and $R_o$, respectively. The annulus has a rectangular cross-section $L \times W$, where $L = R_o - R_i$ is the length of the chamber along the plane of the annulus and $W$ is the chamber width along the optical axis (Fig. S7A). Since the cross-section is long and thin i.e. $L \gg W$, the fluid locally responds to the nearest walls, and therefore it is sufficient to consider the fluid motion at the center of the annulus $R = (R_i + R_o)/2$, along a one dimensional section along the smallest chamber dimension, which is the width $W$ (see Fig. S7A). Also since $R_i, R_o \gg W$, we neglect the effects of the chamber curvature at leading order. At leading order, the effects of gravity also do not contribute to fluid motion and only result in a hydrostatic gradient given by:

$$p_{static}(z) = p_{chamber} + (R_o - z)\rho_f g, \tag{11}$$

where $p_{chamber}$ is the ambient pressure in the chamber, and the height $z$ is measured from the horizontal plane at the 3 O'clock azimuthal location of the circular chamber (Fig. S7A). With these assumptions, the problem for the fluid motion reduces to that of start-up flow in a channel of width $W$, where the fluid motion is purely due to the motion of the channel walls (Fig. S7B). For convenience, we use a coordinate system where the channel walls are at $y = 0,\ W$ (Fig. S7B). Since the chamber is circular, the problem reduces to one where the channel is infinite in extent along the length, therefore making the problem one dimensional (Fig. S7B).

We now solve for the fluid motion in this configuration for a start-up motion of the walls. With the above assumptions, the flow is uni-directional so that $\boldsymbol{u} = u_z(t, y)\boldsymbol{e_z}$ and the dynamic pressure field can be written as $p_{dynamic} = p(t, y)$. The fluid flow equations, therefore, reduce to a diffusion equation for $z$-momentum [9] which is given by:

$$\frac{\partial u_z}{\partial t} = \nu \frac{\partial^2 u_z}{\partial y^2}, \tag{12}$$

with the following initial and boundary conditions:

$$\begin{cases} u_z(t=0,y) = u_0(y) = 0, & \textit{initial conditions} \\ u_z(t,0) = u_z(t,W) = U, & \textit{boundary conditions}. \end{cases} \tag{13}$$

Here $U$ is the velocity of the walls.

Solution of this equation is standard [9], and can be solved by decomposing the solution to a general and particular solution, where the general solution satisfies a homogeneous boundary condition. A standard separation of variables type approach results in the solution for the fluid velocity:

$$u_z(t,y) = U\left[1 - \frac{4}{\pi}\sum_{n=0}^{\infty}\frac{1}{(2n+1)}\exp\left(-\frac{(2n+1)^2\pi^2}{W^2}\nu t\right)\sin\left(\frac{(2n+1)\pi y}{W}\right)\right], \tag{14}$$

and the solution for the pressure is $p_{dynamic} = 0$.

This constitutes the response of the fluid to a step change in velocity of amplitude $U$. Thus one can write down the step response of the fluid to a unitary step function in wall velocity, or equivalently, the impulse response of the fluid to a delta distribution in wall acceleration, as follows:

$$u_\delta(t,y) = \left[1 - \frac{4}{\pi}\sum_{n=0}^{\infty}\frac{1}{(2n+1)}\exp\left(-\frac{(2n+1)^2\pi^2}{W^2}\nu t\right)\sin\left(\frac{(2n+1)\pi y}{W}\right)\right]. \tag{15}$$

Having derived the impulse response of the fluid, we can now derive the response to a general velocity profile of the walls. We use the fact that the governing equations for the fluid motion (Eqs. (12), (13)) constitute a linear, time-invariant (LTI) dynamical system, that is, the fluid velocity can be written as:

$$u_z(t, y) = \mathcal{F}[a(t)] \tag{16}$$

where $a(t)$ is the acceleration profile of the walls, and $\mathcal{F}$ is a LTI operator. Since $\mathcal{F}$ is LTI it obeys the convolution theorem, so that we have:

$$u_z(t, y) = u_z(t = 0, y) + \int_0^t a_{wall}(\tau) u_\delta(t - \tau) d\tau \tag{17}$$

where $u_\delta$ is the impulse response derived in Eq. (15). Using this we can derive the fluid velocity profile for general motion of the walls, as shown in Fig. S7D, for the case of a constant acceleration to a maximum velocity. As seen from Eq. (17), this response depends on the past history of wall motion via a kernel that decays over the viscous time scale $W^2/\nu$.

Using the Eq. (17), We find, as expected, that the maximum shear rate always occurs at the walls ($y = 0, W$). Further, we calculate the time dynamics of the shear rate developed in the fluid, which is marked by an initial growth to temporal maximum value followed by a decay to zero, as the fluid velocity becomes a solid body motion with the walls. It is this spatio-temporal maximum in shear rate, that we use in our characterization of the effects of shear on the orientation of tracked objects, discussed next. Since this maximum shear rate occurs at the walls and is also a temporal maximum, it sets the upper bound for the shear rate experienced by a tracked object anywhere in the channel, at any time during its track. Using the relevant value for the maximum stage acceleration used in our experiments, we find that this maximum in shear rate does not exceed about $1 s^{-1}$, at any point of our tracing.

#### 2.3 Effects of stage and fluid response time on tracking

We now consider the effects of stage and fluid response times on vertical tracking. The fluid response time has been derived in §2.2, and has a viscous lag which scales as $\tau_{visc} \sim W^2/\nu$. The stage response time is determined by the specific motion profile implemented on our

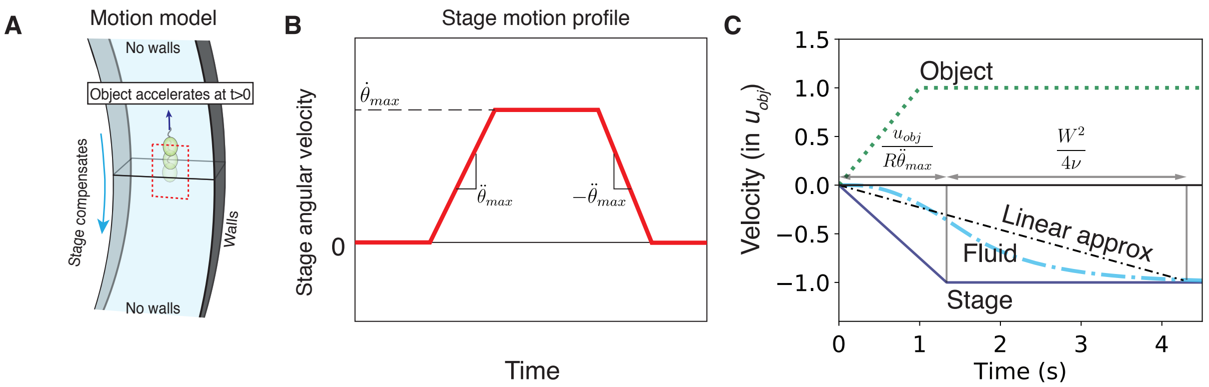

**Figure S8** Effects of stage and fluid response time on tracking. **(A)** Motion model for an organism or object that changes its vertical velocity by $u_{obj}$. To track this motion, the stage compensates by accelerating in the opposite direction at a rate $\ddot{\theta}_{max}$. **(B)** Motion profile implemented on our rotational stage. A change in position, is implemented by acceleration at a constant rate $\ddot{\theta}_{max}$ to a maximum speed $\dot{\theta}_{max}$ and deceleration at a constant rate $-\ddot{\theta}_{max}$. This angular velocity profile maps to a translational velocity of the walls at the object's radial location as $a_{stage} = R\dot{\theta}(t)$. **(C)** Translational velocity vs time of the object (green dotted curve), stage (solid blue curve) and fluid (cyan dash-dotted curve). The fluid velocity lags behind that of the stage due to the viscous delay of $\tau_{visc} \sim W^2/\nu$ in the transfer of momentum. For analytical calculations, this delay can be modelled, without worrying about the actual details of the fluid-velocity time-series (cyan dash-dotted curve) using a linear approximation (black dash-dotted curve), as shown.

rotational stage. This motion profile consists of acceleration at a rate $\ddot{\theta}_{max}$ to a maximum angular velocity of $\dot{\theta}_{max}$, and is given in Fig. S8 B.

We consider a motion model for a tracked object, at the channel center-line, wherein it accelerates and changes its vertical velocity by $u_{obj}$ at an acceleration $a_{obj}$, and the stage compensates by moving in the opposite direction at an acceleration $a_{stage} = R\ddot{\theta}_{max}$, where $R$ is the instantaneous radial location of the object (Fig. S8A). The object's velocity relative to the fluid is:

$$u_{obj|fluid}(t) = \begin{cases} a_{obj}t, & t \leq u_{obj}/a_{obj} \\ u_{obj}, & t > u_{obj}/a_{obj} \end{cases} \tag{18}$$

The stage's velocity relative to the lab is:

$$u_{stage|lab}(t) = \begin{cases} -a_{stage}t, & t \leq u_{obj}/a_{stage} \\ -u_{obj}, & t > u_{obj}/a_{stage}. \end{cases} \tag{19}$$

While we have numerically derived the fluid's velocity profile in §2.2, in order to make analytical progress towards deriving an expression for the tracking error as a function of time, we make a simplifying assumption for the velocity time dynamics of the fluid. We make a linear approximation for the fluid velocity such that the fluid accelerates at a constant rate, but at a rate slower than the stage (Fig. S8C). Therefore while the stage takes a time $\tau_{stage} = u_{obj}/(R\ddot{\theta}_{max})$ to accelerate to the object's velocity, the fluid at the channel center-line takes a time given by $\tau_{stage}+\tau_{visc} = u_{obj}/(R\ddot{\theta}_{max})+W^2/4\nu$, where $W^2/4\nu$ is the time-scale for momentum to diffuse from the channel walls to its center-line. Therefore, the fluid velocity at the center-line can be approximated as:

$$u_{fluid|lab}(t) = \begin{cases} -\frac{u_{obj}}{\tau_{stage}+\tau_{visc}}t, & t \leq \tau_{stage} + \tau_{visc} \\ -u_{obj}, & t > \tau_{stage} + \tau_{visc} \end{cases} \tag{20}$$

Since the Reynolds and Stokes numbers are small for the objects we consider (see §2.1), both fluid and object inertia is negligible, and the object velocity relaxes to that set by the instantaneous fluid drag. Based on this, the object's velocity relative to the lab (or microscope's FOV) is given by:

$$u_{obj|lab}(t) = \begin{cases} a_{obj}t - \frac{u_{obj}}{\tau_{stage}+\tau_{visc}}t, & t \leq u_{obj}/a_{obj} \\ u_{obj} - \frac{u_{obj}}{\tau_{stage}+\tau_{visc}}t, & u_{obj}/a_{obj} < t \leq \tau_{stage} + \tau_{visc} \\ 0, & t > \tau_{stage} + \tau_{visc}. \end{cases} \tag{21}$$

Note that we have made the assumption above that $u_{obj}/a_{obj} < \tau_{stage} + \tau_{visc}$, which is the relevant limit to consider since we are interested in deriving the limits of our tracking method for fast object dynamics. Using this we get the vertical tracking error (displacement of the object relative to the microscope FOV) as:

$$z_{error}(T) = \int_0^T u_{obj|lab}(t)\ dt. \tag{22}$$

Note that it is sufficient to consider $T < \tau_{stage} + \tau_{visc}$, since the velocity at later times is zero. Using Eq. (21), we can write Eq. (22) as:

$$z_{error}(T) = \int_0^{u_{obj}/a_{obj}} \left(a_{obj} - \frac{u_{obj}}{\tau_{stage} + \tau_{visc}}\right) t\ dt + \int_{u_{obj}/a_{obj}}^{T} \left(u_{obj} - \frac{u_{obj}}{\tau_{stage} + \tau_{visc}} t\right)\ dt. \tag{23}$$

To obtain a tracking condition, we require that the vertical tracking error for an object starting at the center of the FOV not cross the FOV, which can be written as the inequality:

$$z_{error}(T) \leq L_{FOV}/2, \tag{24}$$

where $L_{FOV}$ is the vertical extent of the microscope's FOV. Substituting Eq. (23) in Eq. (24), integrating and simplifying, we obtain the following inequality:

$$T^2 - 2(\tau_{stage} + \tau_{visc})T + (\tau_{obj} + L_{FOV}/u_{obj})(\tau_{stage} + \tau_{visc}) \geq 0. \tag{25}$$

To derive the tracking condition, one can use the equality condition for Eq. (25), since that denotes the point when the object reaches the edge of the FOV. Thus the condition for tracking success at all times requires that equation $T^2 - 2(\tau_{stage} + \tau_{visc})T + (\tau_{obj} + L_{FOV}/u_{obj})(\tau_{stage} + \tau_{visc}) = 0$, have no real roots. This gives us the condition for tracking success as:

$$\boxed{\tau_{visc} + \tau_{stage} \leq \frac{L_{FOV}}{u_{obj}} + \tau_{obj}} \tag{26}$$

#### 2.4 Effects of transient shear on object orientation

We now consider the effects of transient shear on the orientation of the tracked objects. This shear is generated since changes to the rotational velocity of the fluidic chamber leads to a transient non-uniform flow profile, which on the scale of a microscale object is a transient shear flow (Fig. S9 A). Thus in the limit $d \ll W$, where $d$ is the object size, the flow is locally given by $\boldsymbol{u} = -\dot{\gamma} y \hat{\boldsymbol{e}}_{\boldsymbol{z}}$, where $\dot{\gamma}$ is the shear rate and $\hat{\boldsymbol{e}}_{\boldsymbol{z}}$ is the unit vector along the z-direction (upward direction). This flow can be decomposed into an extensional and vortical component, where the extensional tensor is given by:

$$\mathbf{E} = \begin{bmatrix} 0 & 0 & 0 \\ 0 & 0 & -\dot{\gamma}/2 \\ 0 & -\dot{\gamma}/2 & 0 \end{bmatrix} \tag{27}$$

and the vorticity is $\boldsymbol{\omega} = \dot{\gamma} \hat{\boldsymbol{e}}_{\boldsymbol{x}}$.

We model the tracked organism as a prolate spheroid with aspect ratio $q$, with a bottom-heavy density distribution which is parameterized by a separation distance $\Delta d$ between the center-of-buoyancy and center-of-mass. The orientation of such a spheroidal, bottom-heavy object can be written using Jeffery's theory [12, 13, 14] as:

$$\dot{\boldsymbol{p}} = \frac{1}{2B}[\hat{\boldsymbol{e}}_{\boldsymbol{z}} - (\hat{\boldsymbol{e}}_{\boldsymbol{z}}.\boldsymbol{p})\boldsymbol{p}] + \frac{1}{2}\boldsymbol{\omega} \wedge \boldsymbol{p} + \beta[\mathbf{E}.\boldsymbol{p} - (\mathbf{E} : \boldsymbol{pp})\boldsymbol{p}], \tag{28}$$

where $B = \mu\alpha_{\perp}/(2g\rho_{obj}\Delta d)$, is a time-scale for reorientation due to gravity, $\mu$ is the dynamic viscosity of the fluid, $\alpha_{\perp}$ is a drag coefficient for rotation of a prolate spheroid [15], $\rho_{obj}$ is the object's mean density, and $\beta = (1 - q^2)/(1 + q^2)$.

Substituting the expressions for the extension tensor and vorticity in Eq. (28), and further simplifying for the case of small angles $\theta$, we have, to leading order in $\theta$:

$$\dot{\theta} = -\frac{\theta}{2B} + \frac{\dot{\gamma}(1 - \beta)}{2}. \tag{29}$$

The first of the terms on the right-hand-side of Eq. (29) represents the orientation stabilizing, gravitactic term, while the second is the destablizing term due to the shear. Each of these terms is additionally the inverse of a time-scale. Thus, to quantify the relative strengths of the terms we can consider the ratio of the two time-scales, namely the gravitactic time-scale $\tau_{grav} = B$, and the shear time scale $\tau_{shear} = [\dot{\gamma}(1-\beta)]^{-1}$. The object's orientation is therefore stable and orientation perturbation due to shear, negligible when:

$$\boxed{\frac{\tau_{shear}}{\tau_{grav}} \gg 1} \tag{30}$$

For the sake of our analysis to determine the operating limits of vertical tracking, we further consider the spatio-temporal maximum in shear rate, so as to include a large safety margin.

#### 2.5 Point spread function of the optical system

Point spread function of the optical system is simulated by Zemax. That the object plane is in water and the presence of a 1.5 $mm$ thick acrylic wall are taken into account (Figure S10). The root-mean-square (RMS) voltage applied to the liquid-lens is set to be 39.5V, which is the offset value used in experiments. Cross section of the point spread function and its maximum intensity projection to the y and z axes are plotted in Figure S10. Lateral and Axial FWHM are 2.8 $\mu m$ and 116 $\mu m$ respectively.

#### 2.6 Liquid-lens chracterization

Relationship between the RMS voltage applied to the liquid-lens and the relative working distance is obtained by imaging a calibration slide mounted on a translation stage and recording the stage micrometer reading when the slide is adjusted to be in focus for different liquid-lens RMS voltages. Linear fit of the data yields a coefficient of 73.8$\mu m/V_{rms}$ (Figure S10).

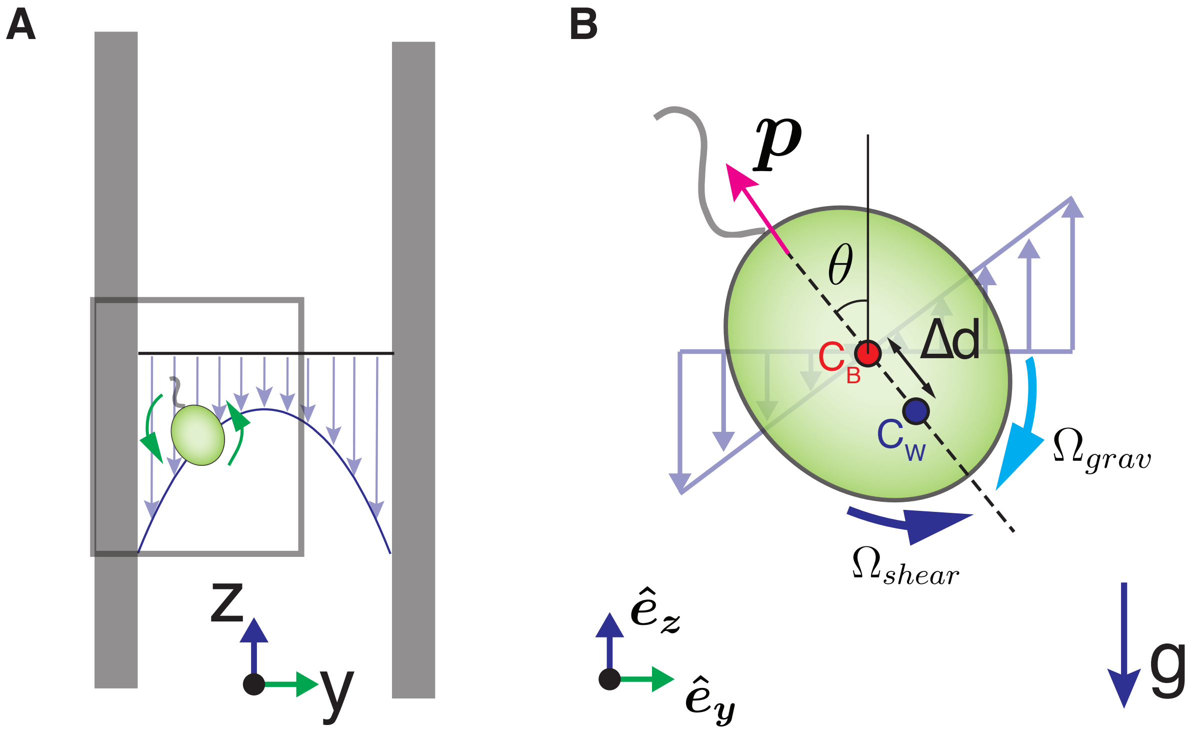

**Figure S9** Effects of transient shear on object orientation. **(A)** Acceleration of the circular fluidic chamber causes a transient shear in the fluid which at the scale of a microscale object is a simple shear flow. **(B)** This simple shear flow can modify the object's natural orientation ($\boldsymbol{p}$) which is set by a bottom-heavy density distribution that tends to align the orientation to the vertical. The bottom-heavy density distribution can be parametrized by a separation $\Delta d$ between the center-of-buoyancy ($C_B$) and the center-of-mass ($C_M$).

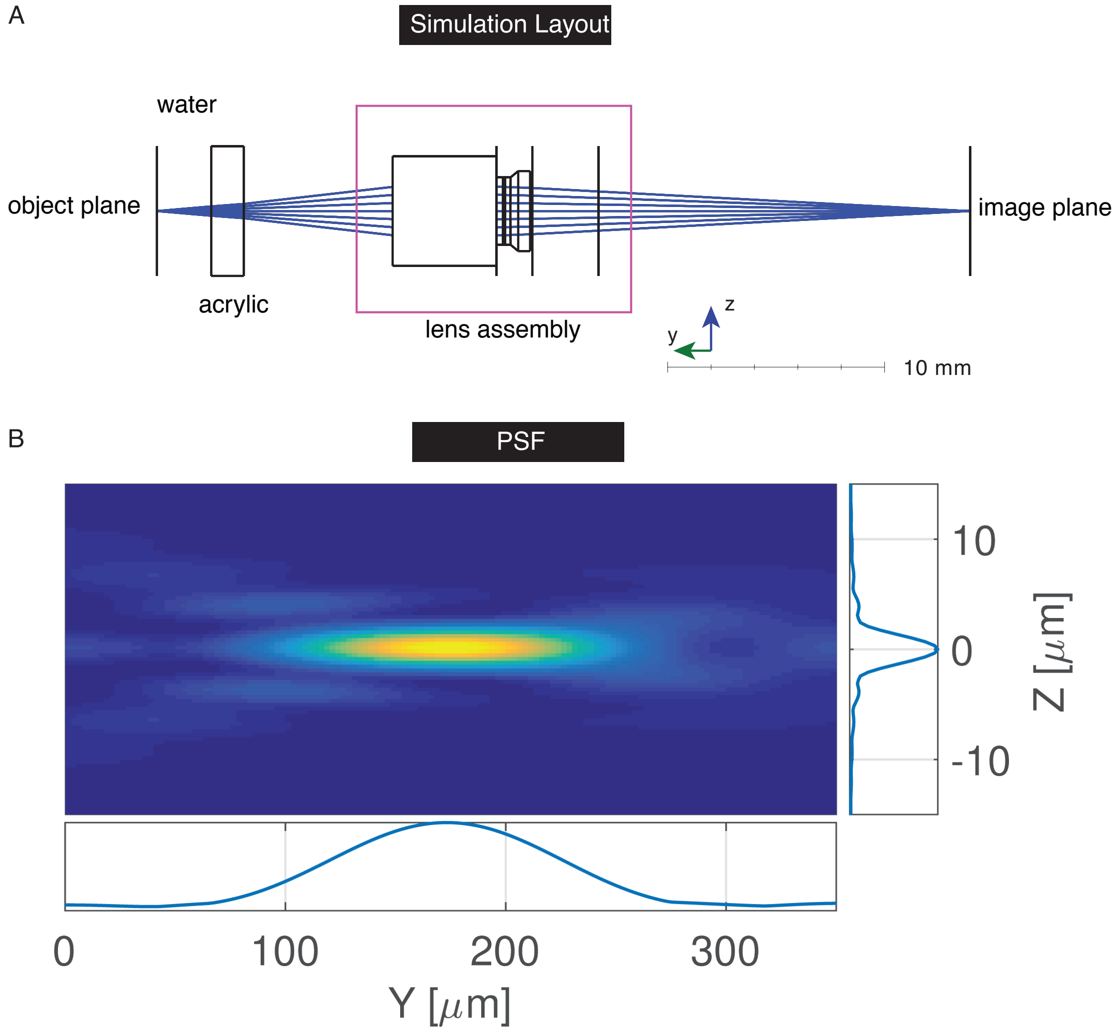

**Figure S10** Point spread function of the optical system. **(A)** Simulation layout. The point spread function is simulated in Zemax. In the simulation a black box model of the liquid-lens assembly provided by the vendor (Corning) is used. The simulation takes into account the presence of acrylic chamber wall and the fluid (assumed to be fresh water with refractive index of 1.33). **(B)** Simulated point Spread Function. Lateral and Axial FWHM are $2.8\ \mu m$ and $116\ \mu m$ respectively.

Frequency response of the liquid-lens is obtained by converting the modulation of optical power (inverse of the focal length) to modulation of beam size of a large diameter laser beam that is focused by the liquid-lens in a fixed plane where the nominal beam size is a few millimeters (Figure S11). A biased photodetector (DET036A, ThorLabs, load resistor not shown) is placed such that part of the beam overlaps with the detector. The voltage output, along with the sine waveform, generated using an Arduino's built-in DAC, that encodes the RMS voltage applied to the liquid-lens, are recorded by an oscilloscope. The frequency of the modulation is swept and the amplitude and relative phase (between the two recorded waveforms) are extracted by fitting sine functions to the recorded oscilloscope traces. The obtained amplitude and phase response are shown in Figure S11. The phase response measured here is used to generate a look-up table that applies a phase correction when using the liquid-lens for our focus tracking method.

| Larva name | Larva type | Days post fertilization |
|---|---|---|
| *S. californicum* | Tornaria | 28 |
| *D. excentricus* | Pluteus | 11 |
| *S. purpuratus* | Early Pluteus | 2 |
| *P. miniata* | Bipinnaria | 12 |
| *P. parvimensis* | Gastrula | 2 |
| *Owenia spp.* | Trocophore | 4 |
| *O. spiculata* | Pluteus | 12 |
| *C. fornicata* | Veliger | 13 |

**Table S1** Marine invertebrate larvae information.

| Bead type | Color code | Size range ($\mu m$) | Density ($g\ cc^{-1}$) |
|---|---|---|---|
| 0 | Green | 212 - 250 | $1.02 \pm 0.005$ |
| 1 | Orange | 212 - 250 | $1.04 \pm 0.005$ |
| 2 | Violet | 212 - 250 | $1.06 \pm 0.005$ |
| 3 | Dark blue | 250 - 300 | $1.08 \pm 0.005$ |
| 4 | Red | 212 - 250 | $1.09 \pm 0.005$ |
| 5 | Blue | 212 - 250 | $1.13 \pm 0.005$ |

**Table S2** Density calibration beads (Cospheric) used for control experiments.

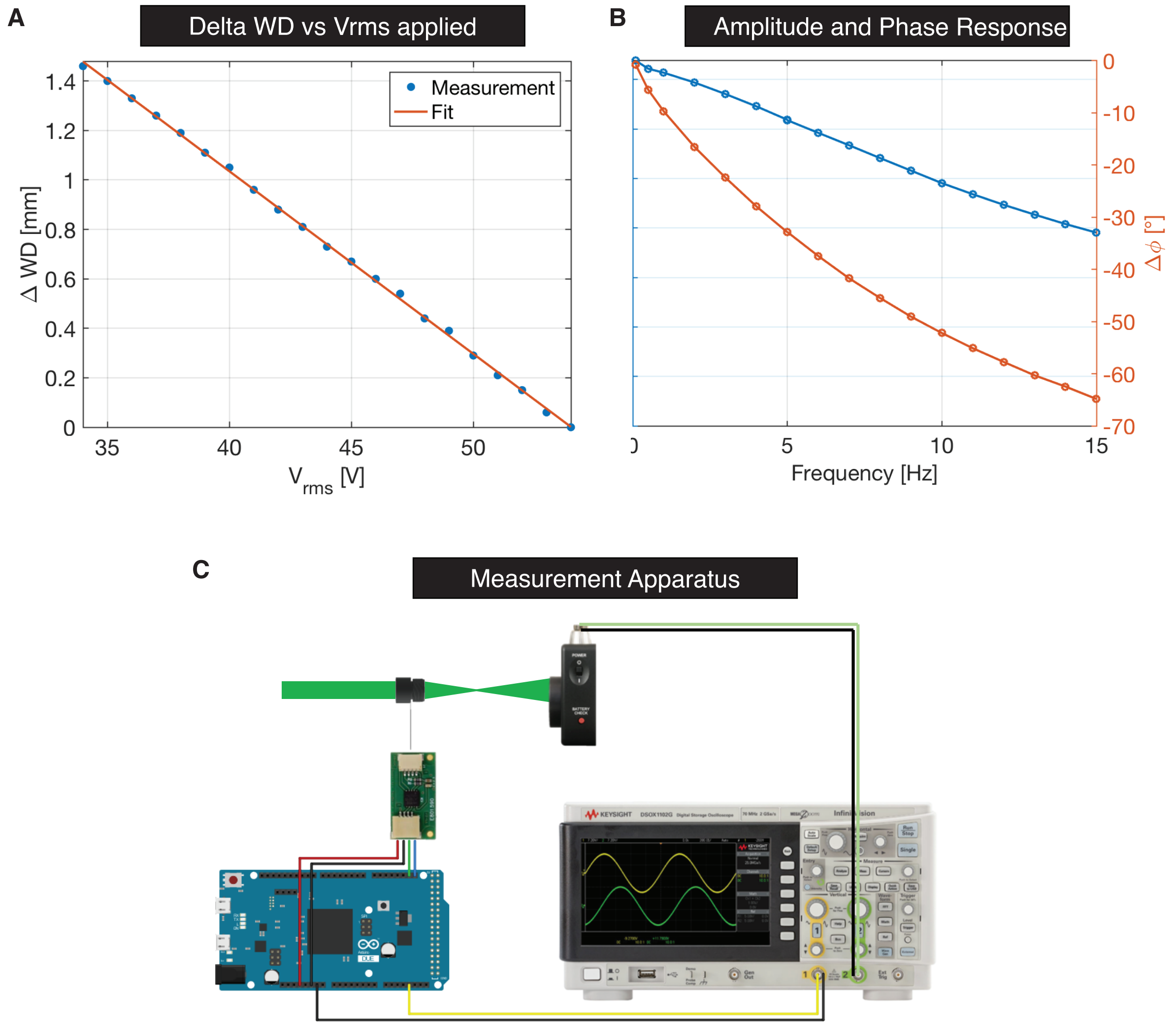

**Figure S11** Liquid-lens Characterization. **(A)** Measured change in working distance vs voltage (root mean squared) applied to the liquid-lens. 20 V change in applied voltage (root mean squared) corresponds to 1.48 mm change in working distance, with coefficient of $73.8\mu m/V_{rms}$. **(B)** Amplitude and phase response of the liquid-lens vs modulation frequency. **(C)** Measurement apparatus for obtaining the amplitude and phase response.

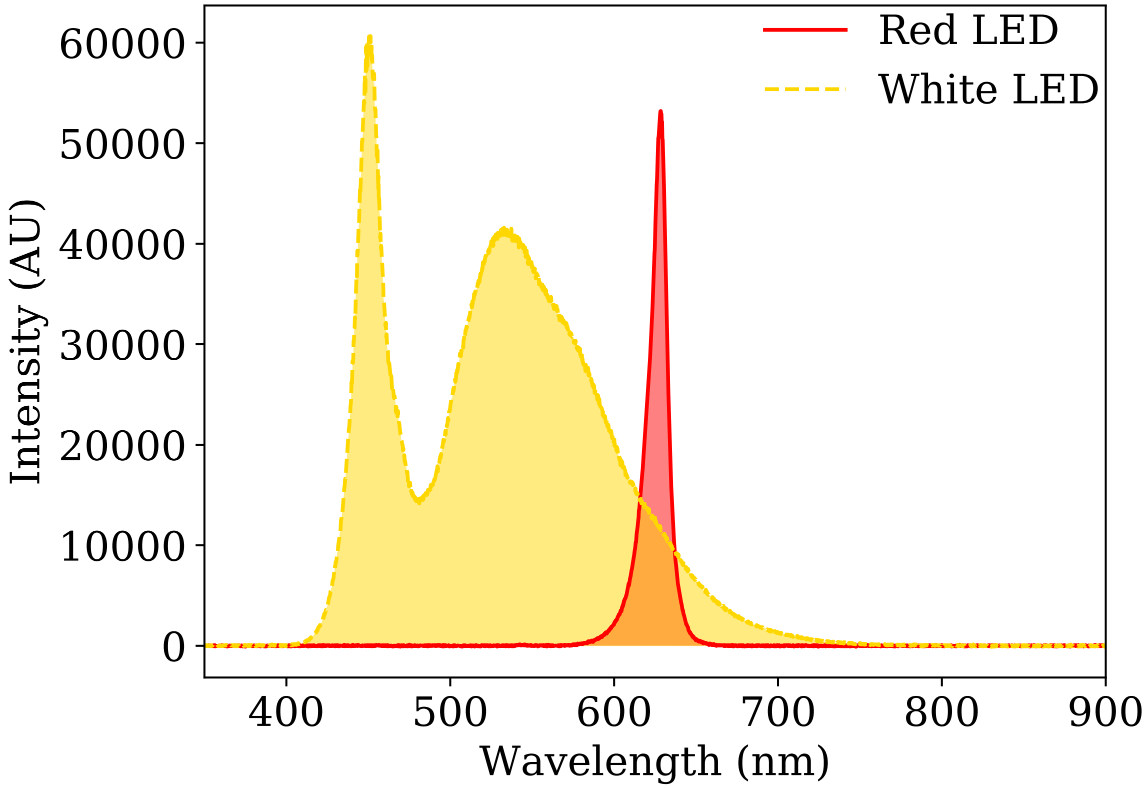

**Figure S12** Measured intensity spectra for the two types of LED illumination used in experiments. The spectrum was measured using an Ocean Optics Spectrometer and a standard desktop computer.

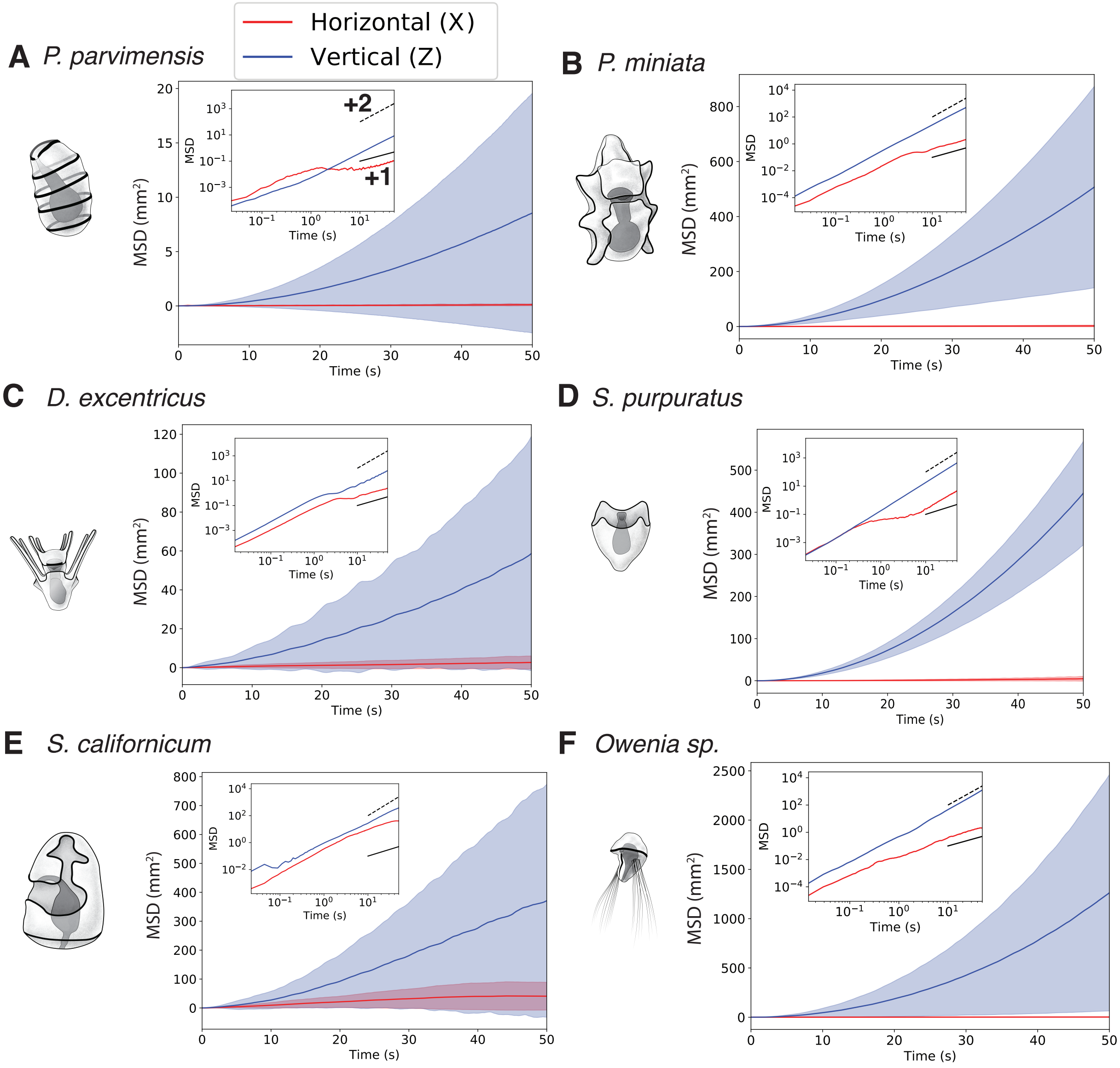

**Figure S13** Measured vertical (z) and horizontal (x) Mean-Squared-Displacements (MSD) vs time for different marine invertebrate larvae. The insets show a log-log plot of MSD vs time, with the slopes corresponding to ballistic (dashed line) and diffusive (solid line) trajectories marked. For all larvae, except those of *S. californicum*, the vertical MSD remains ballistic even for long times, whereas the horizontal MSD becomes diffusive.

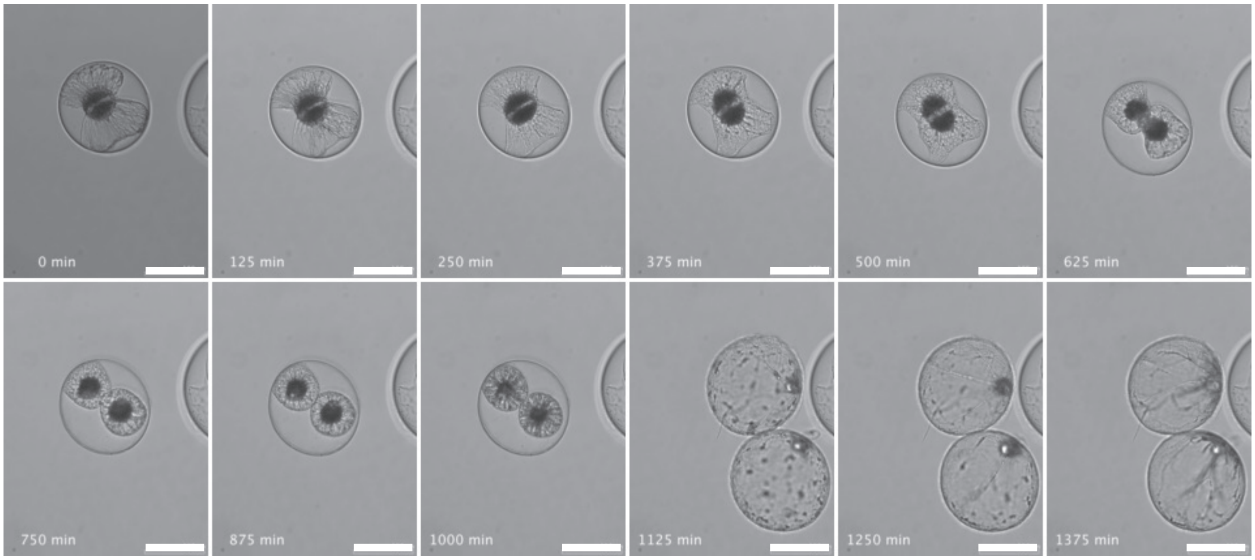

**Figure S14** Control experiment with *P. noctiluca*. Snapshots in time of *P. noctiluca* undergoing cell-division observed under a Nikon Ti2-E inverted microscope using DIC imaging. The rapid expansion in the daughter cells volume post-division is seen.

| PIV Parameter | Parmeter value/range |
|---|---|
| Pixel size | 1.6 $\mu m$ |
| Window size | 64 pixels |
| Search area | 64 pixels |
| Overlap | 32 pixels |
| Frame rate | $>$ 30 Hz |
| Bead size | 2 or 3 $\mu m$ |
| Bead concentration | 0.05% by volume |

**Table S3** Parameters used for Particle Image Velocimetry (PIV).

### 3 Supplementary Movie Captions

**Movie 1** Setup and procedure for a vertical tracking experiment.

**Movie 2** Multi-scale measurements of sedimenting particles: (1) Interactions in Sedimenting Spheres , (2)Rods, and (3)Microscale transport processes in coupled sedimentation-dissolution.

**Movie 3** Tracking of sedimenting marine detritus and concurrent measurement of microscale transport.

**Movie 4** A comparative study of marine invertebrate larvae behavior.

**Movie 5** Multiscale tracking of freely swimming *P. miniata* (bat-star) larvae reveal behavioral transitions that regulate depth and enable feeding.

**Movie 6** Diel behavior of Polychaete larvae measured at the scale of individual organisms.

**Movie 7** Measuring of behavior of plankton in depth-patterned virtual environments: Volvox colony response to depth-dependent changes in light intensity.

**Movie 8** Tracking single cells: Dynamic sinking behavior in marine diatoms.

**Movie 9** Tracking single cells: Measuring cell division and associated changes in sinking rates in the dinoflagellate *Pyrocystis noctiluca*.

### References

[1] A. Lukežič, T. Vojí, L. ČehovinZajc, J. Matas, M. Kristan, *International Journal of Computer Vision* **126**, 671 (2018).

[2] Z. Zhu, *et al.*, *Lecture Notes in Computer Science (including subseries Lecture Notes in Artificial Intelligence and Lecture Notes in Bioinformatics)* **11213 LNCS**, 103 (2018).

[3] C. F. Batten, D. M. Holburn, B. C. Breton, N. H. M. Caldwell, *Scanning* **23**, 112 (2001).

[4] C. F. Batten, *Mphill thesis, University of Cambridge* (2000).

[5] M. F. Strathmann, *Reproduction and development of marine invertebrates of the northern Pacific coast: data and methods for the study of eggs, embryos, and larvae* (University of Washington Press, 2017).

[6] K. Drescher, R. E. Goldstein, N. Michel, M. Polin, I. Tuval, *Physical Review Letters* **105**, 1 (2010).

[7] A. Liberzon, *et al.*, OpenPIV-Python.

[8] W. Gilpin, V. N. Prakash, M. Prakash, *The Journal of Experimental Biology* **220**, 3411 (2017).

[9] L. G. Leal, *Advanced transport phenomena: fluid mechanics and convective transport processes*, vol. 7 (Cambridge University Press, 2007).

[10] E. A. Salzen, *Experimental Cell Research* **12**, 615 (1957).

[11] S. Kim, S. J. Karrila, *Microhydrodynamics: principles and selected applications* (Courier Corporation, 2013).

[12] G. B. Jeffery, *Proceedings of the Royal Society of London. Series A* **102**, 161 (1922).

[13] J. S. Guasto, R. Rusconi, R. Stocker, *Annual Review of Fluid Mechanics* **44**, 373 (2012).

[14] T. Pedley, *Annual Review of Fluid Mechanics* **24**, 313 (1992).

[15] H. C. Berg, *Random walks in biology* (Princeton University Press, 1993).
